# Brief synaptic inhibition persistently interrupts firing of fast-spiking interneurons

**DOI:** 10.1101/2022.08.02.502477

**Authors:** Simon Chamberland, Erica R. Nebet, Manuel Valero, Monica Hanani, Robert Egger, Samantha B. Larsen, Katherine W. Eyring, György Buzsáki, Richard W. Tsien

## Abstract

Neurons perform input-output operations that integrate synaptic inputs with intrinsic electrical properties, operations generally constrained by the brevity of synaptic events. Here we report that sustained firing of CA1 hippocampal fast-spiking parvalbumin-expressing interneurons (PV-INs) can be persistently interrupted for up to several hundred milliseconds following brief GABA_A_R-mediated inhibition *in vitro* and *in vivo*. A single presynaptic neuron could interrupt PV-INs firing, occasionally with a single action potential (AP), and reliably with AP bursts. Experiments and computational modeling revealed that the persistent interruption of firing maintains neurons in a depolarized, quiescent state through a cell-autonomous mechanism. Strikingly, interrupted PV-INs are highly responsive to Schaffer collateral inputs. The persistent interruption of firing provides a disinhibitory circuit mechanism favoring spike generation in CA1 pyramidal cells. Overall, our results demonstrate that neuronal silencing can far outlast brief synaptic inhibition owing to well-tuned interplay between neurotransmitter release and postsynaptic membrane dynamics, a phenomenon impacting microcircuit function.

## Introduction

Synaptic excitation and inhibition drive or prevent action potential (AP) firing to gate information transfer in neuronal circuits. In cortical networks, synaptic inhibition is mediated by functionally heterogeneous GABAergic interneurons (INs) (Freund and Buzsaki, 1996; Klausberger and Somogyi, 2008; Pelkey et al., 2017). Among these, fast-spiking parvalbumin (PV)-expressing INs (PV-INs) are recognized as powerful modulators of neuronal network activity (Cardin et al., 2009; Sohal et al., 2009; Stark et al., 2013) and behavior (Donato et al., 2013; Kuhlman et al., 2013; McKenna et al., 2020). Although PV-INs represent a minority of neurons in the hippocampus, the synaptic inhibition they provide contributes to network oscillations such as those associated with memory formation (Royer et al., 2012; Amilhon et al., 2015). This ability is thought to be supported by their extensive axonal arborization (Sik et al., 1995), the powerful inhibitory connections they form on their postsynaptic targets (Bartos et al., 2002; Hefft and Jonas, 2005), and their intrinsic biophysical properties. In response to depolarization, PV-INs generate non-accommodating bouts of high-frequency APs (Kawaguchi et al., 1987), a phenotype enabled by the combined activity of Na_v_1.1, Na_V_1.6, and K_v_3-family channels which rapidly depolarize and repolarize the membrane (Martina et al., 1998; Rudy and McBain, 2001; Lorincz and Nusser, 2008; Hu and Jonas, 2014). *In vivo*, this rapid AP discharge is phase-locked to ongoing network activity with bouts of firing interspersed with periods of relative silence (Klausberger et al., 2003; Klausberger et al., 2004; Klausberger and Somogyi, 2008). Therefore, PV-INs appear integral to the coordination of neuronal network activity.

Synaptic inhibition arising from other INs has been demonstrated as a powerful factor constraining the activity of GABAergic INs (Cobb et al., 1997; Gulyas et al., 1999; Chamberland and Topolnik, 2012; Tyan et al., 2014). This action of inhibitory neurons onto other GABAergic neurons leads to a net disinhibitory effect in the neuronal network, enabling the passage, processing, and storage of information. This idea is well-exemplified by findings showing that inhibition of PV-INs is involved in associative fear learning (Letzkus et al., 2011; Wolff et al., 2014). At the network level, it has long been suggested that interconnected populations of INs may entrain ensembles of pyramidal cells (Buzsaki et al., 1983; Lytton and Sejnowski, 1991). Reciprocal connectivity among PV-INs contributes to the emergence of network activity such as gamma oscillations (Wang and Buzsaki, 1996; Bartos et al., 2002; Bartos et al., 2007). In the CA1 hippocampus, subpopulations of vasoactive intestinal peptide (VIP)-expressing INs have long been recognized as disinhibitory neurons and somatostatin-expressing (SST-INs) are known to synapse onto PV-INs (Acsady et al., 1996; Lovett-Barron et al., 2012). While key in understanding circuit function, the inhibitory synaptic wiring diagram to PV-INs is incomplete.

Synaptic inhibition operates by either hyperpolarizing the membrane potential or shunting incoming excitatory inputs. The release properties of the presynaptic neuron and the postsynaptic receptor subtypes constrain the duration of inhibition. Considering that PV-INs are assembled in densely interconnected inhibitory networks (Sik et al., 1995; Gulyas et al., 1999; Acsady et al., 2000; Bartos et al., 2001) and have a resting membrane potential relatively depolarized compared to other interneuron types (Gentet et al., 2010; Tricoire et al., 2011; Yu et al., 2016), GABAergic synapses are poised to profoundly affect PV-INs. Although the classical view holds that neuronal input-output transformation happens on the timescale of synaptic activity, evidence from multiple brain regions shows that neuronal firing can be maintained or emerge following stimulus termination (Kiehn and Eken, 1998; Egorov et al., 2002; Shu et al., 2003a; Fransen et al., 2006; Sheffield et al., 2011; Cui and Strowbridge, 2019). The ability of neurons and neuronal networks to generate episodes of persistent activity may enable information to be retained for periods of time exceeding the original stimulus (Durstewitz et al., 2000; Egorov et al., 2002; Shu et al., 2003a) and thus provide a physiological substrate for operations such as working memory. Yet, whether and how synaptic inhibition can switch neurons between different firing states is largely unexplored.

Here, we discovered a mechanism based on the interplay between inhibitory synaptic transmission and intrinsic membrane properties that prolongs the silent period exhibited by PV-INs in response to minimal synaptic inhibition, a phenomenon we term persistent interruption of firing. Our analysis reveals that the persistent interruption of firing results from an interplay between a D-type K^+^ current and a Na^+^ current that work together to keep PV-INs quiescent yet hyperresponsive. The interruption of firing is a disinhibitory mechanism for AP firing in CA1 pyramidal neurons.

## Results

### Synaptic inhibition interrupts firing of fast-spiking interneurons

Synaptic inhibition hyperpolarizes the membrane potential relative to the AP threshold, silencing the neuron. The duration of postsynaptic neuron silencing is thought to result from the combination of presynaptic release properties and kinetics of postsynaptic receptor activation. PV-INs fire APs at high frequency upon depolarization. Yet, how other interneurons affect PV-IN activity remain generally obscure.

To understand how synaptic inhibition controls PV-INs activity, we performed experiments in acute hippocampal slices prepared from P17 – P30 animals. PV-INs were depolarized with suprathreshold current sufficient to evoke their characteristically fast and sustained firing (Fig. 1A-B). Synaptic inhibition was elicited through optogenetic stimulation of somatostatin-expressing interneurons (SST-INs) by using *Sst;;Ai32* transgenic mice. We chose this approach because SST-INs form synaptic contacts on PV-INs, while SST- and PV-INs represent generally non-overlapping populations of INs (Freund and Buzsaki, 1996; Jinno and Kosaka, 2000; Harris et al., 2018; Udakis et al., 2020). Optogenetic stimulation of SST-INs afferent with a 20 ms light pulse during sustained PV-INs firing strongly suppressed subsequent firing (Fig. 1A-C), leaving the neuron in a non-firing depolarized state (average membrane potential of −36.4 ± 1.1 mV during the interruption compared to −66.6 ± 0.6 mV at rest, n = 29; p < 0.001). We termed this phenomenon a persistent interruption of firing (also referred to as an interruption for brevity), in contrast to the brief silencing expected for an IPSP. The same optogenetic inhibition applied when recording from depolarized CA1 pyramidal neurons (CA1-PYR) revealed no such persistent interruption (Fig. S1A-C). The likelihood of observing inhibitory currents in PV-INs and CA1-PYR upon optogenetic stimulation of SST-interneurons was not different and IPSCs displayed similar properties (Fig. S1D-E), resulting in similar IPSP amplitudes (PV-INs: 3.82 ± 0.56 mV, n = 29; CA1-PYRs: 5.57 ± 1.32 mV; n = 9; p = 0.34, Mann-Whitney U test). These results show that persistent interruption of firing is a selective mechanism for powerfully controlling PV-IN activity.

**Figure 1:**
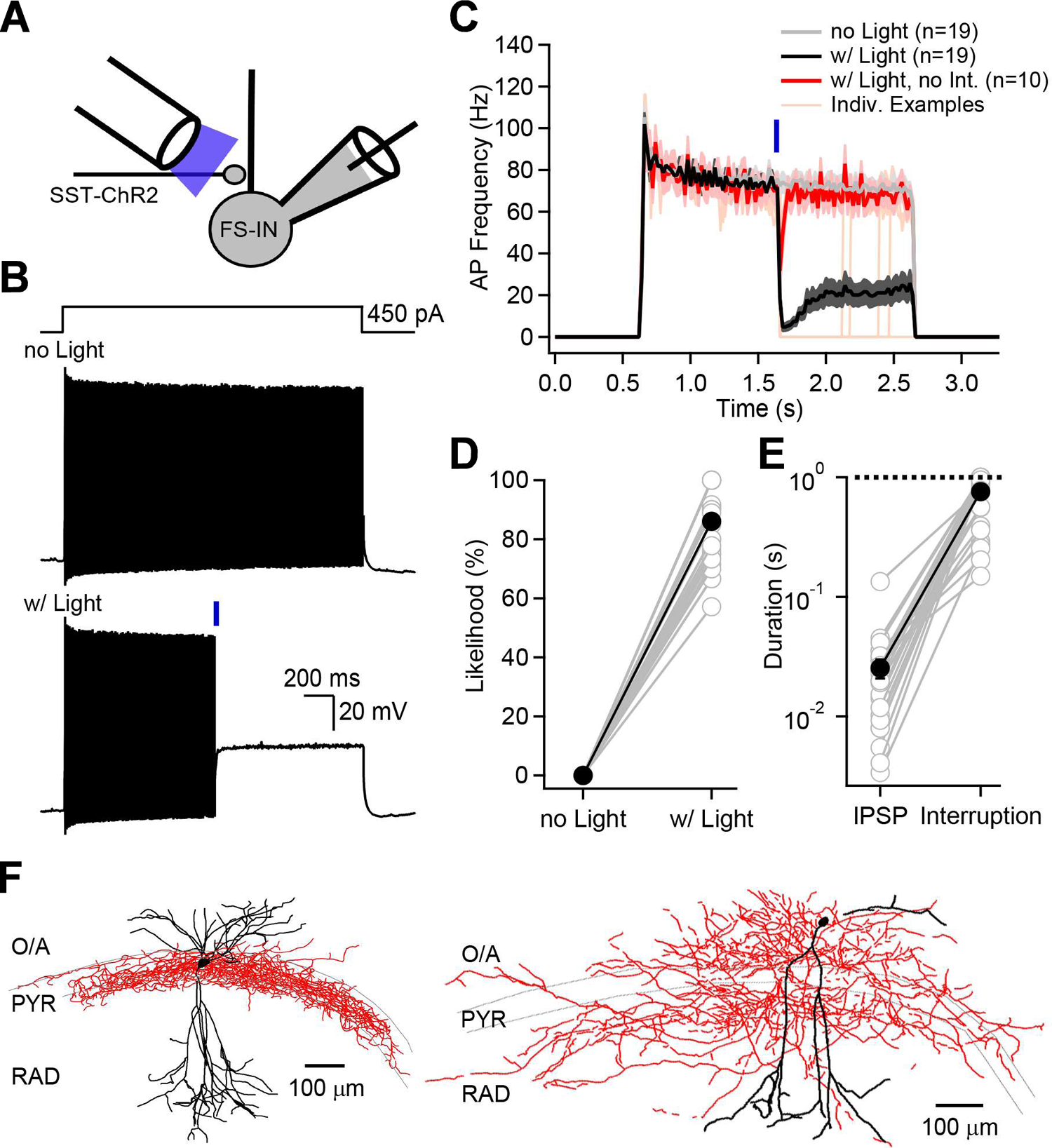
Synaptic inhibition persistently interrupts firing of PV-Ins. **A**, Scheme showing the recording configuration. **B**, PV-INs were depolarized with a square current injection to elicit firing. In absence of optogenetic stimulation, PV-INs demonstrate a classical fast and non-adapting spiking phenotype. Brief optogenetic stimulation generated an IPSP followed by an interruption of firing. **C**, Summary data of AP firing frequency as a function of time for experiments shown in B, in absence (light gray) or presence of optogenetic stimulation (black). The red trace shows the average of traces when no firing interruption was induced, and the orange traces show individual trials as examples. **D**, Likelihood of observing a firing interruption for all PV-INs sampled. **E**, Duration in seconds of the IPSP compared to the silence period imposed by the firing interruption. The dashed line represents the duration of the depolarizing step, therefore capping the possible interruption duration value at 1 s. **F**, Neurolucida anatomical reconstructions of PV-INs recorded and filled with biocytin. The dendrites are shown in black and the axon is shown in red.

Persistent interruption of firing was observed with a high reliability in response to optogenetic intervention (86.1 ± 2.4%, n = 29; Fig. 1C-D). In trials where the interruption of firing was not induced, PV-INs firing rapidly recovered (Fig. 1C, red trace) and the silence duration was similar to that observed in pyramidal cells (Fig. S1C). In interleaved trials, we observed that PV-INs maintained their firing in the absence of optogenetic intervention (Fig. 1C and Fig. S2A, B).

Additionally, most PV-INs (17/29 neurons) could resume normal firing, in which case the initial firing frequency was fully recovered (before interruption: 76.3 ± 4.2 Hz; after interruption: 72.3 ± 4 Hz, n = 17; p = 0.37, Mann-Whitney U test). Measuring the duration of the interruption revealed that the silencing lasted on average 757 ± 56 ms (n = 29), approximately 30-fold longer than the full duration of the IPSP (25.4 ± 4.5 ms, n = 29, Fig. 1E). There was no significant correlation across neurons between the initial firing rate and the likelihood of observing an interruption (Pearson correlation: r = 0.1, p = 0.59; n = 29; Fig. S1F). The firing interruption was observed at all temperatures tested, with the baseline firing of PV-IN reaching an average of >300 Hz at 31.3 ± 0.9°C (Fig. S1G-I). On the other hand, increasing the depolarizing pulse amplitude by 37.2 ± 3.6% of the suprathreshold current value (from 301.9 ± 46.9 pA to 413.4 ± 66.9 pA, n = 3 neurons) prevented the persistent interruption of firing, enabling the firing to resume right after the IPSP. Next, the duration of the optogenetic stimulation was decreased to explore the synaptic determinants generating the interruption (Fig. S2C-K). Brief light pulses (2 ms) generated fewer APs in SST-expressing INs (Fig. S2C-E) but still generated interruptions of similar durations, albeit with a lower likelihood (Fig. S2G-I). The briefer (2 ms) optogenetic stimulation evoked IPSPs of similar amplitude, but of shorter duration (Fig. S2J-K). Increasing the intracellular Cl^-^ concentration to shift E_Cl_ closer to physiological values reported in PV-INs (−52 mV and −64 mV) (Vida et al., 2006; Otsu et al., 2020) slightly decreased the likelihood of observing the persistent interruption of firing (E_Cl_ −52 mV: 73.3 ± 8.3%, n = 7; E_Cl_ −69 mV: 86.1 ± 2.4%, n = 29; p = 0.14, Mann-Whitney U test; Fig. S2L-M). Thus, like hyperpolarizing inhibition, even shunting inhibition (Vida et al., 2006) sufficed.

Post-hoc anatomical reconstructions of 23 PV-INs and cluster analysis based on axonal distribution allowed us to separate the recorded neurons into two groups; 1) perisomatic-targeting cells with an axon ramifying in *stratum pyramidale* (Fig. 1F and Fig. S3A) and, 2) dendrite-targeting cells with an axon innervating *strata oriens* and/or *radiatum* (Fig. 1F and Fig. S3B). All perisomatic-targeting (7/7) and dendrite-targeting (16/16) neurons demonstrated persistent firing interruptions, with similar likelihood (perisomatic-targeting: 87.1 ± 3.6 %; dendrite-targeting: 85.5 ± 4.0 %, p = 1.0, Mann-Whitney U test) and duration (perisomatic-targeting: 888.6 ± 39.9 ms; dendrite-targeting: 665.3 ± 84.6 ms, p = 0.18, Mann-Whitney U test). This indicates the robustness and the ubiquity of this phenomenon amongst different types of PV-INs. Therefore, the persistent interruption of firing is a novel mechanism greatly prolonging the inhibition interval generated by GABAergic input to two subtypes of PV-INs.

### A single action potential at a unitary connection suffices to interrupt firing

Optogenetic experiments revealed that even small IPSPs can interrupt PV-IN firing for hundreds of milliseconds. PV-INs receive synaptic inhibition from multiple sources, including PV-INs themselves (Chamberland and Topolnik, 2012). However, which presynaptic neurons must be recruited and how many presynaptic APs are required to interrupt PV-INs firing remains unclear.

Paired recordings were performed to determine the presynaptic activity required to interrupt PV-INs firing (Fig. 2A). We found that firing from a single presynaptic partner was sufficient to interrupt firing in most synaptically-connected pairs interrogated (14 out of 16 connected pairs, Fig. 2B-D). Furthermore, a single AP evoked by a single presynaptic partner was sufficient to interrupt PV-INs in a subset of connected pairs (5 out of 11 connected pairs), but with a low likelihood (Fig. 2B, D). Delivering a brief burst of 5 APs at 100 Hz, a physiological pattern of activity for interneurons (Klausberger et al., 2003; Klausberger et al., 2004), was considerably more efficient at interrupting the firing (1 AP: 4.42 ± 2.26 %; 5APs: 36.74 ± 7 %; n = 10; p < 0.001, Mann-Whitney U test), with the interruption likelihood plateauing for yet more APs (10 APs: 41.61 ± 7.62 %; Fig. 2C-D). The firing interruption initiated by a single presynaptic partner, while of lower likelihood than in optogenetic experiments, demonstrated nearly identical duration (paired recordings: 739 ± 68 ms, n = 12; optogenetics:757 ± 56 ms, n = 29; p = 0.67, Mann-Whitney U test). Neurolucida reconstructions revealed that in all cases, the postsynaptic neurons had anatomical features consistent with PV-INs (Fig. S3A-E). We observed that for presynaptic PV-INs, 6/10 neurons projected their axon in the dendritic layers, while 4/10 neurons innervated the perisomatic region (Fig. 2A and S3D). When the presynaptic partner was an SST-IN, the axon was found in dendritic layers in 4/4 neurons.

**Figure 2:**
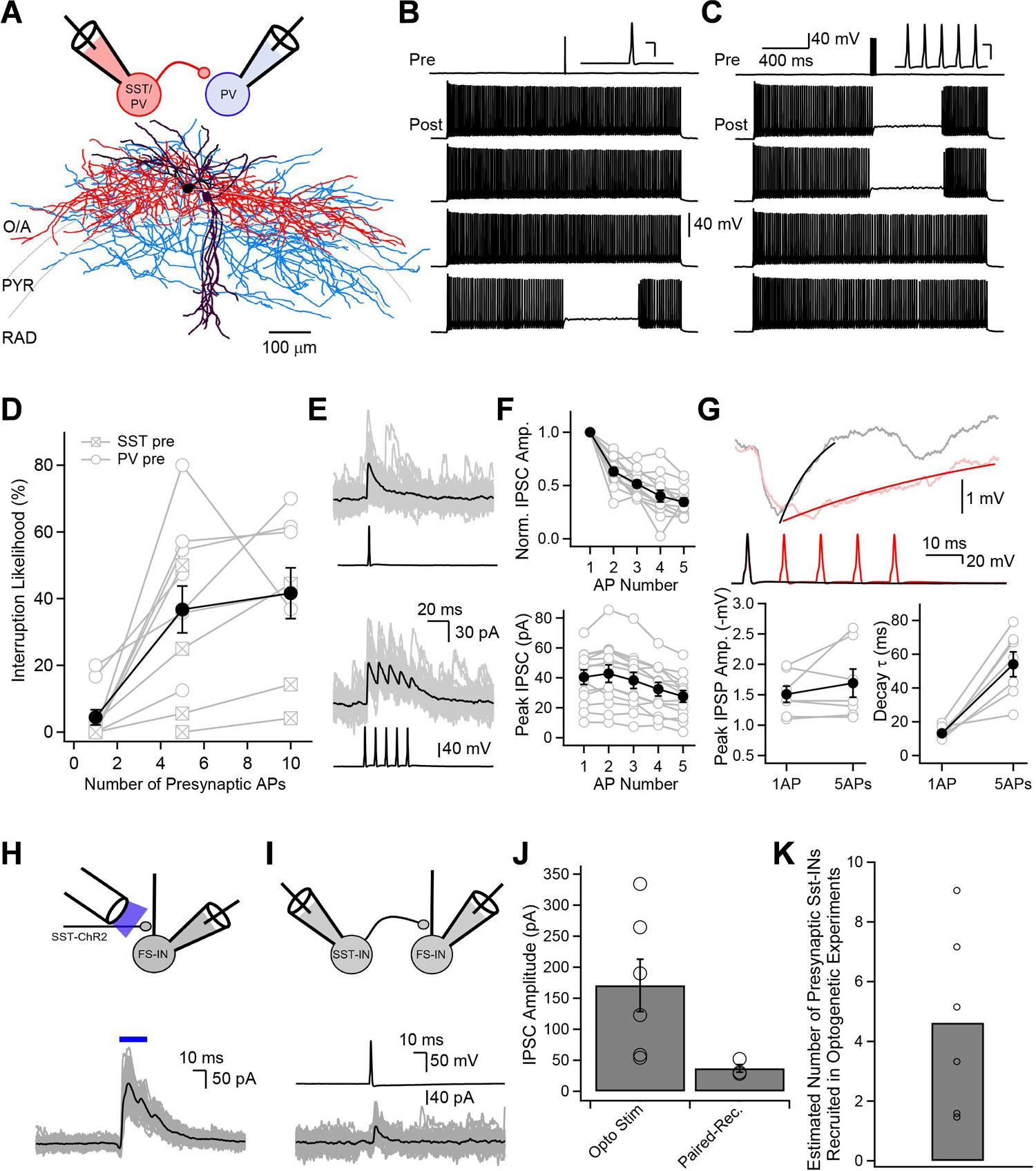
A single presynaptic interneuron can interrupt PV-INs firing. **A**, Recording configuration and post hoc Neurolucida reconstruction of a synaptically-connected pair of interneurons. The dendrites of the presynaptic interneuron are shown in black and its axon is shown in red. The dendrites of the postsynaptic neurons are shown in purple and the axon is in blue. **B**, Current-clamp recordings in a pair of synaptically-connected interneurons. A single AP evoked by current injection in the presynaptic cell is sufficient to interrupt post-synaptic firing. Four consecutive epochs are shown for the postsynaptic interneurons. **C**, Same pair as in B, with five APs evoked at 100 Hz in the presynaptic cell. Insets scale bar for B and C: 40 mV vertical, 5 ms horizontal. **D**, Summary graph showing the firing interruption likelihood as a function of the number of presynaptic APs. The five and ten AP bursts were delivered at 100 Hz. The amplitude of the depolarizing current injection in the postsynaptic PV-INs was 255 ± 22.55 pA (n = 11). Presynaptic PV- and SST-INs had similar likelihood to interrupt firing when five APs were evoked (n = 7 and n = 4, respectively; p = 0.11). **E**, Voltage-clamp recordings performed at 0 mV in the postsynaptic neurons reveal large IPSCs for single and five APs-evoked bursts. Black traces are the average of 50 consecutive sweeps shown in gray. **F**, top, Normalized IPSC amplitude as a function of stimulus number showing short-term depression. IPSC amplitude was measured from the trough to the peak. bottom, Absolute peak amplitude of the IPSC burst measured from baseline for all AP-evoked IPSC reveals a relatively efficient summation, with the amplitude declining gradually due to the short-term depression. **G**, Current-clamp recordings of single- and five APs-evoked IPSP. Black and red traces are the average of 3 consecutive sweeps. The maximal peak amplitude of the five APs-evoked IPSP is similar to a single AP-evoked IPSP, however the decay kinetics are greatly increased by five APs bursts. IPSCs measured in optogenetic experiments (**H**), in paired-recordings (**I**) and summary graph of IPSC amplitudes (**J**). **K**, Estimation of the number of SST-INs synapsing onto a single PV-IN.

To better understand the mechanisms controlling the interruption, we next analyzed the underlying currents evoked at unitary synaptic connections. Single AP firing reliably generated large amplitude IPSCs in postsynaptic PV-INs (Fig. 2E), characteristics consistent with previous reports (Bartos and Elgueta, 2012). We then analyzed the short-term dynamics of IPSCs evoked by brief trains of presynaptic APs (5 APs at 100 Hz). We observed that bursts of IPSCs demonstrated significant short-term depression (1^st^ IPSC: 40.45 ± 4.72 pA; 5^th^ IPSC: 13 ± 1.67 pA; p < 0.0001, n = 13; Fig. 2E-F) but summated efficiently, such that the absolute peak amplitude of the burst evoked IPSC was maintained for the first two APs and then declined (Fig. 2E-F). Given the short-term depression, we next asked why train-evoked APs were more likely to interrupt firing. Analysis of the resulting IPSP waveform during subthreshold depolarization revealed that the peak amplitude was similar between 1 AP and 5 APs-bursts (1 AP: 1.51 ± 0.12 mV; 5 APs: 1.69 ± 0.22 mV; p = 0.3, n = 7; Fig. 2G), while the decay kinetics were strikingly slowed (1 AP: *τ* = 13.18 ± 1.33 ms; 5 APs: *τ* = 54.05 ± 6.84 ms, p < 0.01, n = 7; Fig. 2G), prolonging the return to the depolarized membrane potential. These results demonstrate that while a single AP from a unitary synaptic connection is sufficient to trigger persistent interruption of PV-IN firing, brief bursts of APs, generating extended inhibition, are significantly more efficient.

It was clear that the interruption likelihood was lower in paired recordings than in optogenetic experiments (Fig. 2D compared to Fig. 1D). We compared the currents evoked by optogenetic stimulation and paired-recordings to estimate how many SST-INs contribute to the total inhibitory current required to reliably trigger firing interruptions. Voltage-clamp recordings in fast-spiking interneurons revealed that optogenetic stimulation of SST-INs evoked large IPSCs of 170.6 +/- 42.3 pA (n = 6 neurons; Fig. 2H-J). In contrast, the unitary IPSC amplitude observed in paired-recordings was 36.9 +/- 6.2 pA (n = 3 synaptically connected pairs; Fig. 2I-J). By comparing the total synaptic drive in each case, we estimated that an average of 4 – 5 SST-INs innervate a single fast-spiking interneuron (Fig. 2K). Therefore, coordinated activity from multiple presynaptic interneurons raises efficiency of interrupting PV-INs.

### Fast-spiking interneurons in vivo can remain silent for an extended duration following brief synaptic inhibition

Our results indicate that PV-INs can be silenced for long periods in response to brief optogenetic activation of inhibitory afferents in acute hippocampal slices. We next explored whether long silent periods indicative of persistent interruption of firing could be induced by synaptic inhibition in the intact brain.

For *in vivo* tests, we combined multisite silicon probe electrophysiological recordings with optogenetic stimulation in behaving *Sst;;Ai32* mice (Fig. 3A) (Valero et al., 2021). In addition to SST-INs, identified by their responsiveness to blue light, other neuronal types were categorized based on their AP waveform and discharge rate (Fig. 3B-D). For example, narrow-waveform interneurons (NW-INs), classified by brief spike duration and rapid rise time (Fig. 3C-D), were identified as putative PV-INs as previously documented (Henze et al., 2000). The optogenetic stimulation of SST-INs silenced PV-INs for intervals extending beyond the blue light stimulus in most trials (Fig. 3E, 9 typical NW-INs shown, each with 1500 trials). The duration of silencing varied across trials, but in all cells a subset of ranked trials reached the maximal duration sampled (0.6 s; Fig. 3E). By averaging trials in the lowest (0-10^th^ percentile), middle deciles (45-55^th^ percentile) and highest deciles (90-100^th^ percentile) across cells, we confirmed that long-lasting inhibition of PV-INs was a phenomenon consistent across all PV-INs sampled (Fig. 3F). On average, middle deciles trials demonstrated that the silent period consistently outlasted the optogenetic stimulation (Fig. 3F). Conversely, averaging trials in the lowest percentiles revealed that PV-INs can also recover their firing rapidly following an inhibitory event, in which case the silence duration was mostly limited to duration of the optogenetic stimulation (Fig. 3F). This finding is consistent with our *in vitro* observations: failure to induce the persistent interruption of firing resulted in only brief silences (Fig. 1C, red trace).

**Figure 3:**
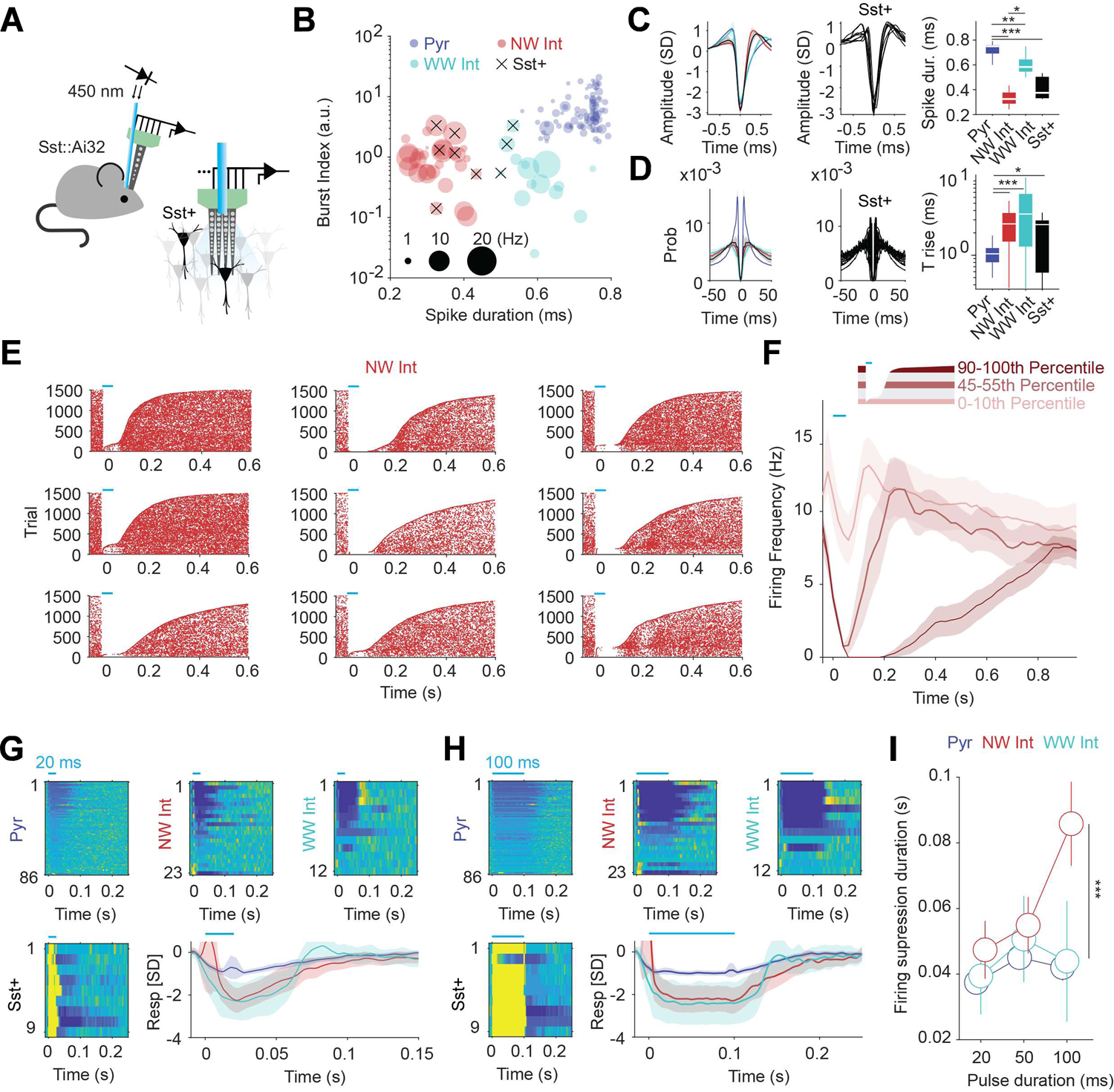
PV-INs silencing persists following optogenetic stimulation *in vivo*. **A**, Schematic of the recording configuration showing combined multisite silicon probe and optical fiber in the CA1 hippocampus of an *Sst;;Ai32* mouse. Animals from the same strains were used for *in vivo* and *in vitro* experiments. **B**, Burst index as a function of spike duration for all neurons sampled (n = 130 units) reveals distinct neuron populations identified as narrow-waveform interneurons (red), pyramidal cells (blue), wide-waveform interneurons (teal) and somatostatin-positive light-sensitive interneurons (black, n = 9 cells). **C**, Average spike waveform for neural population identified in B (left), including all somatostatin-positive light-sensitive interneurons (middle), and through-to-peak spike duration (right). **D**, Same as in C for firing auto-correlograms and rise time to peak. **E**, AP raster plots of 9 narrow-waveform INs during 1500 optogenetic stimulation trials for each cell. Trials are ranked by silencing duration induced by 50 ms optogenetic stimulation. **F**, Summary graph for delay to recovery of spiking for all narrow-waveform INs sampled, showing the averages of trials for the lowest, middle, and highest percentiles across neurons. **G**, **H**, Optogenetic stimulation for 20 ms (G) or 100 ms (H) in other cell types (pyramidal, wide-waveform INs and Sst-expressing INs) results in briefer silencing duration than in narrow-waveform INs. Warmer colors correspond to higher firing rates. **I**, Delay to recovery of spiking as a function of the optogenetic stimulation duration for all cell types. Narrow-waveform INs are silenced by optogenetic stimulation of SST-INs for longer on average of all trials than other cell types. * p < 0.05; ** p < 0.01; *** p < 0.001 for all statistical tests on the data presented.

We next compared the effect of optogenetically-induced inhibition of PV-INs with that of pyramidal cells and wide-waveform interneurons (WW-INs). On average, the silence duration was significantly longer for PV-INs than for other cell types (Fig. 3G-H). This observation is counter-intuitive; given their generally high baseline firing rate *in vivo* (NW: 6.14 ± 3.89 Hz, PYR: 1.02 ± 0.64 Hz, WW: 5.21 ± 4.53 Hz (mean ±SD); NW vs. PYR: p < 0.0001; NW vs. WW: p = 0.4495; ANOVA followed by posthoc Tukey-Kramer), PV-INs would logically be expected to recover their firing faster after synaptic inhibition. Yet, PV-INs were silenced for a consistently longer period than all other neuronal subtypes for all optogenetic stimulus durations sampled (Fig. 3I). This difference was starkest for the 100 ms light pulse duration, consistent with our *in vitro* optogenetic experiments (Fig. S2H-I) and paired recordings (Fig. 2D) showing that a train of stimuli are more likely to induce an interruption of firing and engender a longer silence. Thus, our *in vivo* observations that PV-INs can remain silent for extended periods following optogenetic activation of GABAergic afferents are consistent with our *in vitro* findings in demonstrating similar dependence on cell type and intensity of PV-IN inhibition.

### GABA_A_ receptor blockade prevents and postsynaptic membrane hyperpolarization reproduces the interruption

To understand why the duration of the interruption of firing is variable both *in vitro* and *in vivo*, and why the *in vivo* interruptions are generally briefer, we next set out to examine the underlying biophysical mechanisms controlling the interruption of firing. What are the pre- and postsynaptic events required to persistently interrupt PV-INs? Both optogenetic stimulation and unitary presynaptic neuron firing in paired recordings generated postsynaptic IPSPs, likely mediated by GABA_A_ receptors. Alternatively, non-classical neurotransmission could contribute to the firing interruption through slow postsynaptic inhibition, shunting effects, or sustained release.

Accordingly, we proceeded to dissect the synaptic requirements of the persistent firing interruption, using the optogenetic approach in acute slices from *Sst;;Ai32* mice due to higher throughput and efficiency. Blockade of GABA_A_ receptors with bicuculline fully prevented the persistent interruption of firing in all neurons tested (control: 92.7 ± 4.5 % chance of firing interruption; bicuculline: 0 % chance of firing interruption, n = 6, Fig. 4A, C). This observation does not exclude the possibility of synergistic action of a slow-acting neurotransmitter signaling through G-protein coupled receptors. To address this, we confirmed the presence of persistent interruption of firing following 24 hours of pertussis-toxin treatment to prevent G_i/o_ signaling (n = 3, 85.5 ± 6.7 % chance of firing interruption, Fig. S4A-C) (Eyring et al., 2020). To further test the sufficiency of GABA_A_R signaling, we directly blocked GABA_B_ receptors. GABA_B_R inhibition (2 μM CGP-55845, denoted as CGP) had no effect on the interruption of firing, while subsequent application of bicuculline completely prevented the interruption in the same neurons (n = 8; Fig. S4D-E). Together, these results indicate that presynaptic GABA release and activation of postsynaptic GABA_A_Rs are key steps mediating the persistent interruption of firing.

**Figure 4:**
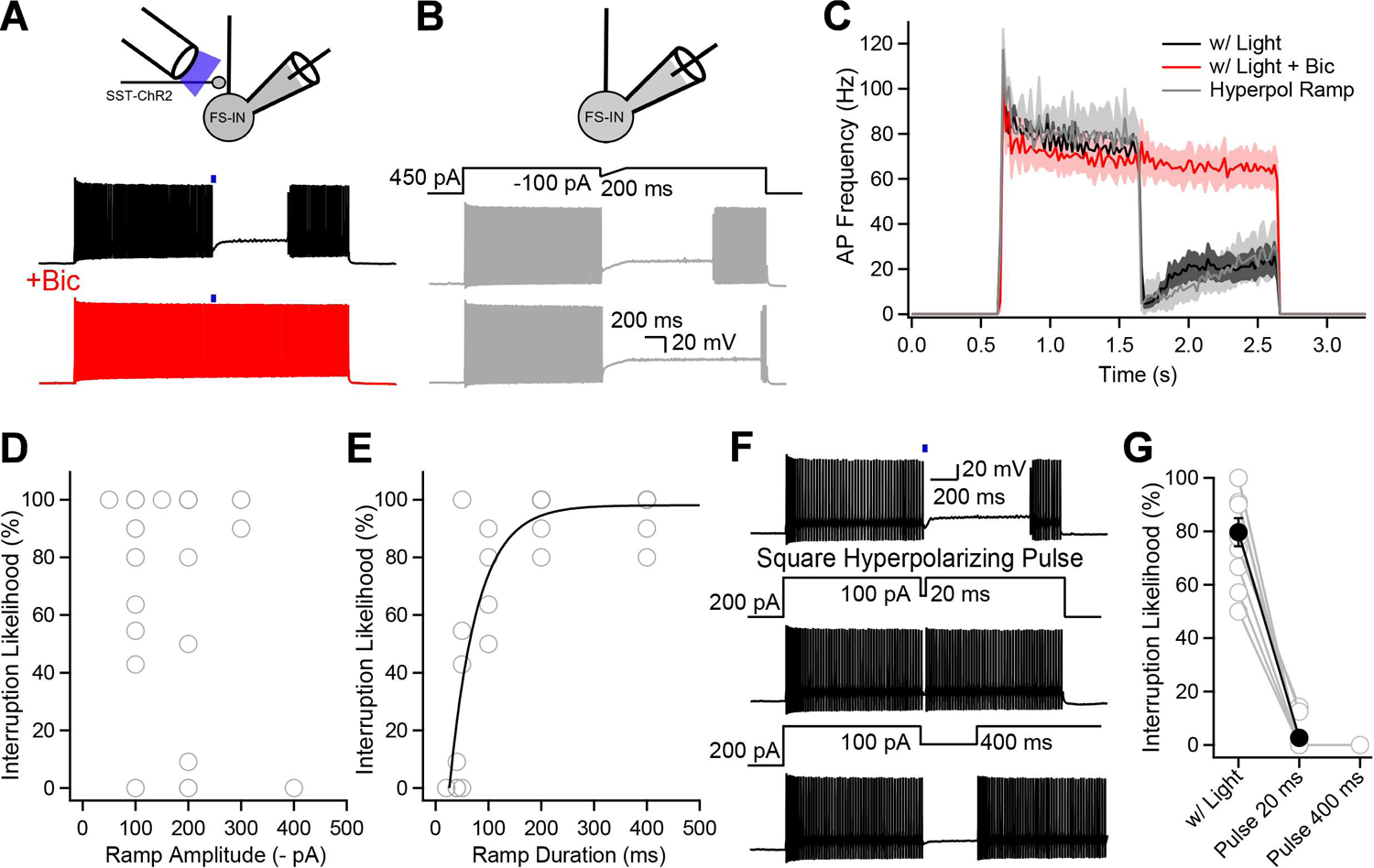
Postsynaptic membrane hyperpolarization through GABA_A_ receptor activation or current injections interrupts PV-INs firing. **A**, Application of the GABA_A_ antagonist bicuculline (10 μM) abolishes the optogenetically-induced interruption of firing. **B**, Hyperpolarizing current injection in PV-INs reliably interrupts their firing. **C**, Summary graph showing the AP frequency as a function of time for optogenetically-evoked stimulation before (black) and after bicuculline application (red). Data is also shown for hyperpolarization-induced interruption, revealing no difference between interruptions evoked by optogenetic stimulation or current injection. **D**, Interruption likelihood as a function of the hyperpolarizing ramp amplitude reveals no significant correlation. **E**, Interruption likelihood plotted as a function of the ramp duration reveals that a slower membrane re-depolarization is more likely to interrupt firing. **F**, A square hyperpolarizing pulse of 20 or 400 ms with a fast membrane re-depolarization fails to interrupt firing. **G**, Interruption likelihood for paired experiments performed with optogenetic stimulation or square hyperpolarizing pulses.

Second, we addressed the postsynaptic factors downstream of GABA_A_R activation that mediate the firing interruption. This was a logical step because PV-INs maintained their stuttering firing pattern despite GABA_A_R and GABA_B_R blockade (n = 7/8 neurons tested in presence of CGP+Bic, Fig. S4F). As GABA_A_R blockade abolished the interruption and GABA_A_R activation elicited clear membrane hyperpolarization in all recordings, we wondered whether mimicking an IPSP waveform with hyperpolarizing current injection might also interrupt PV-IN firing. The current waveform was reduced to two minimal parameters: 1) an instantaneous step to minimal current amplitude, and 2) a ramp recovery to the initial steady current (Fig. 4B). We found that injecting this ramp waveform caused a persistent interruption of firing, with a duration similar to the interruption of firing caused by optogenetic inhibition (ramp: 766.1 ± 109.5 ms; optogenetic: 914.9 ± 39.7 ms; p = 0.23; n = 5, paired t-test, Fig. 4B, C). This result supports the idea that membrane hyperpolarization is sufficient to interrupt firing and shows that the interruption of firing can be induced independently of synaptic transmission. We took advantage of this finding to dissect the key parameters involved in the interruption of firing, varying either the amplitude or the duration of the ramp re-depolarization. While there was no correlation between the interruption likelihood and the peak hyperpolarization amplitude (Pearson correlation: r = −0.15, p = 0.45; n = 8 neurons; Fig. 4D), there was a clear correlation between the interruption likelihood and the ramp duration (Pearson correlation: r = 0.68, p < 0.0001; n = 8 neurons; Fig. 4E). Thus, slower recovery from hyperpolarization engenders a higher likelihood of firing interruption with optogenetic stimulation, paired recordings and with hyperpolarizing ramp currents (Fig. S2H-K and Fig. 2G). In the most extreme case of rapid recovery from hyperpolarization, a square hyperpolarizing pulse almost never interrupted firing (optogenetics: 79.7 ± 5.3 % chance of firing interruption; square hyperpolarizing pulse: 2.7 ± 1.7 % chance of firing interruption; n = 10, Fig. 4F-G). Therefore, these results show that postsynaptic membrane hyperpolarization alone is sufficient to interrupt PV-IN firing, a phenomenon dependent on the speed of recovery from hyperpolarization.

These results suggested that PV-INs firing interruption can be mediated by any presynaptic interneuron subtype if the synaptic inhibition is sufficiently strong and slowly decaying. Vasoactive intestinal peptide-expressing interneurons (VIP-INs) have been shown to preferentially synapse on other INs in the CA1 region of the hippocampus and mediate disinhibition (Acsady et al., 1996; Chamberland et al., 2010; Chamberland and Topolnik, 2012; Tyan et al., 2014; Francavilla et al., 2018; Turi et al., 2019). Therefore, we next tested whether activation of VIP-INs could interrupt PV-INs firing in the *Vip;;Ai32* mouse model (Fig. S4G-I). Optogenetic activation of VIP-INs was generally insufficient to trigger the persistent interruption of firing (interruption likelihood: 2.79 ± 1.5%; n = 15; Fig. S4H-I). To validate these results, we investigated the connectivity of VIP-INs. Given that VIP-INs were shown in paired recordings to target mostly oriens-lacunosum moleculare (OLM) INs in the CA1 *stratum oriens* (Tyan et al., 2014; Francavilla et al., 2018) which are distinct from PV-INs studied here, we decided to use OLM-INs as a positive control. In sequential recordings of neighboring INs from the same slices, we recorded IPSCs from PV-INs and regular-spiking interneurons with I_h_ (classical electrophysiological properties associated with OLMs) in response to optogenetic activation of VIP-INs (Fig. S4J-L). Intriguingly, we found that optogenetic stimulation of VIP-INs produced consistently small IPSCs in PV-INs (19.4 ± 3.2 pA; n = 12) but more than 5-fold larger IPSCs in neighboring OLM-like INs (116.2 ± 16.4 pA; n = 12; p < 0.001; Fig. S4L). This confirms the preferential innervation of OLM-INs by VIP-INs and indicates that VIP-INs only weakly innervate PV-INs in the CA1 hippocampus, explaining their observed inability to impact sustained firing of PV-INs.

### K_v_1 blockade prevents firing interruption

The interruption of firing is initiated by membrane hyperpolarization followed by slow re-depolarization. During the interruption, PV-INs are maintained in a non-spiking but depolarized state. In principle, neurons can be silenced if their membrane potential is kept more hyperpolarized than the AP threshold, or if the membrane potential is depolarized to the point where sodium channels are inactivated, resulting in non-excitability.

Firing interruptions induced during optogenetic stimulation, paired recordings and direct postsynaptic current injection showed some consistent features revealing of the state of excitability: upon resumption of firing, the first AP had a more depolarized take-off potential (pre-int: −35.5 ± 1.11 mV, post-int: −32.19 ± 1.17 mV; p< 0.001; n = 14), a slower maximal dV/dt (pre-int: 164.96 ± 6.25 mV/ms, post-int: 119.31 ± 8.78 mV/ms; p<0.001; n =14), and a smaller amplitude (pre-int: 63.43 ± 1.89 mV, post-int: 51.89 ± 2.89 mV; p< 0.001; n = 14; Fig. 5A-C). The subsequent APs possessed identical characteristics to the last AP before the interruption (values for 2^nd^ AP post-int: take-off potential: −36.43 ± 1.19 mV; p = 0.16; n =14; maximal dV/dt: 158.22 ± 7.97; p=0.15; n = 14; amplitude: 63.47 ± 2.83 mV; p = 0.98; n = 14; Fig. 5A-C). These features are consistent with decreased sodium channel availability during the first spike but not later spikes following the depolarization. Together, these observations suggest that interrupted neurons are maintained in a depolarized quiescent state but not with complete depolarization block, as firing can ultimately resume.

**Figure 5:**
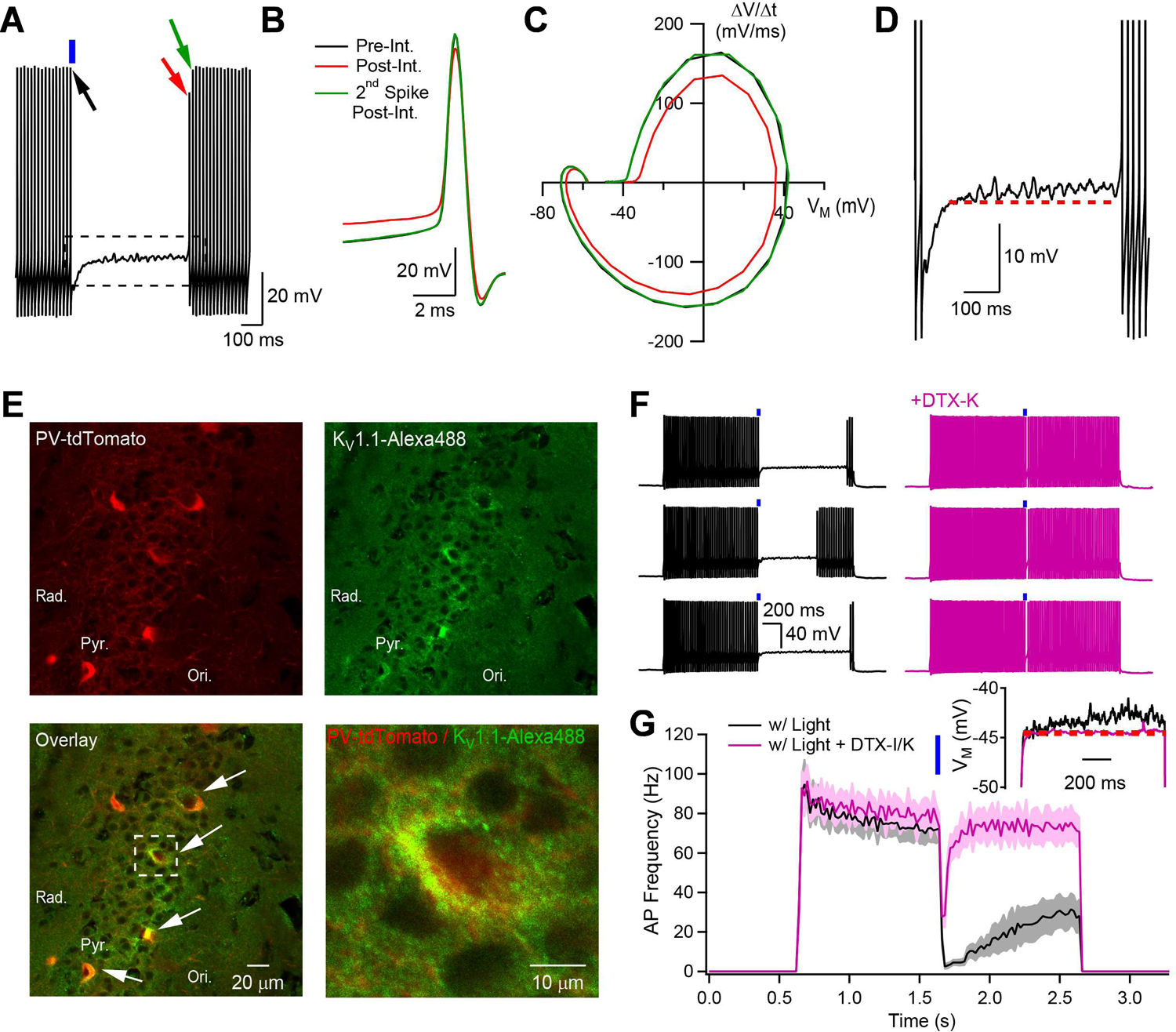
K_v_1.1 is required to interrupt firing. **A**, Current-clamp recording from a PV-IN showing membrane potential dynamics upon firing resumption. **B**, Zoomed-in data from A, showing the APs indicated by arrows. The first AP demonstrates a more depolarized take-off potential and a smaller amplitude. **C**, Phase plot for the three APs shown in B. **D**, During the firing interruption, the membrane potential demonstrates subthreshold oscillations and is gradually depolarized. **E**, Immunohistochemistry experiments reveal that K_v_1.1 is expressed in PV-expressing CA1 interneurons in the regions bordering pyramidal cell layer. White arrows indicate four PV-INs with strong K_V_1.1 correlation at the somatic level. **F**, Optogenetically-induced firing interruption before (black) and after (purple) DTX-K bath application (three consecutive epochs are shown for both control and DTX-K). **G**, AP frequency as a function of time for experiments performed in control and in presence of DTX-K or DTX-I. Inset shows that DTX-K mostly prevents the gradual membrane depolarization upon depolarizing current injection.

To clarify the postsynaptic currents initiating and maintaining the quiescent state, we examined the membrane potential during the firing interruption. All recordings demonstrated a small, slow, and progressive membrane depolarization with a slope averaging 1.71 ± 0.32 mV/s (n = 28) during the interrupted phase (Fig. 5D). Such gradual depolarization could result from a small net inward current arising from a persistent sodium current (I_NaP_), or the gradual inactivation of an outward current. D-type potassium currents (I_D_) mediated by the K_v_1 channel family conduct a gradually inactivating outward current. For molecular constraints, we analyzed published data on the expression of K_v_1 in PV-INs (Cembrowski et al., 2016), which showed relatively higher levels of *Kcna1* and *Kcna2* transcripts compared to moderate levels of *Kcna3* and *Kcna6,* while *Kcna4* and *Kcna5* were mostly undetected (summarized in Fig. S5A). Using immunohistochemistry, we found that hippocampal CA1 PV-INs in the vicinity of stratum pyramidale expressed K_v_1.1 (Fig. 5E). The K_v_1.1 immunoreactivity was prominent in the somatic region of PV-INs. We next tested the involvement of K_v_1.1 in the firing interruption through selective pharmacological blockade with dendrotoxin-I (DTX-I, 50 nM), which blocks K_v_1.1, K_v_1.2 and K_v_1.6, or dendrotoxin-K (DTX-K, 50 nM), which selectively blocks K_v_1.1. Application of either DTX-I or DTX-K prevented the persistent interruption of firing, limiting the quiescent period to that observed in trials where the interruption failed to be induced (Fig. 5F-G, Fig. 1C). We observed that DTX application simultaneously caused a general increase in the AP firing rate in response to depolarization (control: 77 ± 6.2 Hz; DTX-I/K: 88.6 ± 6.9 Hz; n = 9; p < 0.001, Fig. S5B), driven in part by a significant hyperpolarizing shift of the AP take-off potential in DTX without changes in other parameters (Fig. S5C, see legend). To avoid potential confounding effects of increased firing following DTX application, we re-adjusted the depolarizing step amplitude to keep the AP frequency similar to that observed in control condition (Fig. 5F-G). Even so, the likelihood of observing a persistent interruption of firing was strikingly reduced, from 92.5 ± 2.1% to 15.5 ± 4.9% in dendrotoxin (n = 9, including 3/9 neurons in which the interruption was fully abolished; p < 0.0001). Consistent with a proposed role of I_D_ in maintaining the quiescent state during the interruption, the slow and sustained depolarization observed during a square subthreshold depolarizing pulse was virtually absent following DTX-I/K treatment (control: 0.87 ± 0.2 mV/s; DTX-I/K: 0.13 ± 0.06 mV/s; p<0.05; n = 5; Fig. 5G, inset).

K_v_1 channels are formed as heteromultimers incorporating four pore-forming subunits that can include K_v_1.2 and K_v_1.3. To approach the possible roles of K_v_1.2 and K_v_1.3, we exposed neurons to K-Conotoxin RIIIK to block K_v_1.2-containing channels and observed that this decreased the likelihood of firing interruption (control: 100 ± 0%; K-Conotoxin RIIIK: 46.8 ± 15.1%, n = 5; p < 0.05; Mann-Whitney U test; Fig. S5D). On the other hand, bath application of Agitoxin-2, which selectively blocks K_v_1.3, had no significant effect on the firing interruption likelihood (control: 100 ± 0%; Agitoxin-2: 91.3 ± 4.3%, n = 6; p = 0.07; Mann-Whitney U test; Fig. S5E). Overall, these results indicate that K_v_1.1-containing channels are key mediators of the firing interruption and that some of these channels might also contain K_v_1.2 subunits.

### I_D_ and I_NaP_ cooperate to create a stable point in membrane potential

Our evidence for a crucial role for K_v_1.1 in the persistent interruption of firing fits with previous findings that inactivation of K_v_1.1 current powerfully influences AP timing in fast-spiking INs in neocortex (Goldberg et al., 2008). In order to understand how K_v_1.1-mediated currents contribute to the interruption of firing, we next dissected the membrane currents evoked by membrane depolarization and interrogated the membrane dynamics during the interruption.

To assess the full current-voltage relationship over a wide range of membrane potentials, PV-INs held under voltage-clamp were gradually depolarized with a slow ramp from −60 mV to 0 mV over 2 s (Fig. 6A). Exposure to TTX, aimed at pharmacological blockade of Na^+^ currents, was followed by application of DTX-K, directed toward blockade of K_v_1.1 and the corresponding traces were then subtracted to reveal the TTX-sensitive (mostly Na^+^, referred to as ‘I_TTX-s_’ for brevity) and DTX-K-sensitive currents (mostly K_v_1.1, referred to as ‘I_DTX-s_’ for brevity) (Fig. 6B). We next plotted the I-V relationships of the inward I_TTX-s_ and outward I_DTX-s_ (Fig. 6C) to focus on current components with strongly non-linear properties. The summation of I_TTX-s_ and I_DTX-s_ (blue trace, Fig. 6C) suggested the possibility of a stable point in membrane potential where both I_TTX-s_ and I_DTX-s_ exhibit sizable amplitudes, but the net current crosses the zero-current axis with positive slope. There are two logical predictions that can be validated experimentally: first, the membrane conductance should be elevated during the interruption; second, small perturbations to the membrane potential should be followed by a rebound back to the previous level. To determine if these predictions held true, we injected small (50 pA) hyperpolarizing current pulses during the interrupted phase and 2 s later, after recovery of the resting membrane potential (Fig. 6D). Indeed, the input resistance was decreased during the interruption relative to its basal value, consistent with the predicted elevation in membrane conductance (Fig. 6E-F). Furthermore, cessation of the current injection was followed by rebound depolarization (or rebound hyperpolarization, not shown) back to the original quiescent level, indicative of an underlying stable point (Fig. 6E). These results show that the persistent interruption of firing is a shunted quiescent state, corresponding to a stable point in membrane potential, as would be expected for interplay between elevated I_DTx-s_ and I_TTX-s_ as dominant components, acting in opposition.

**Figure 6:**
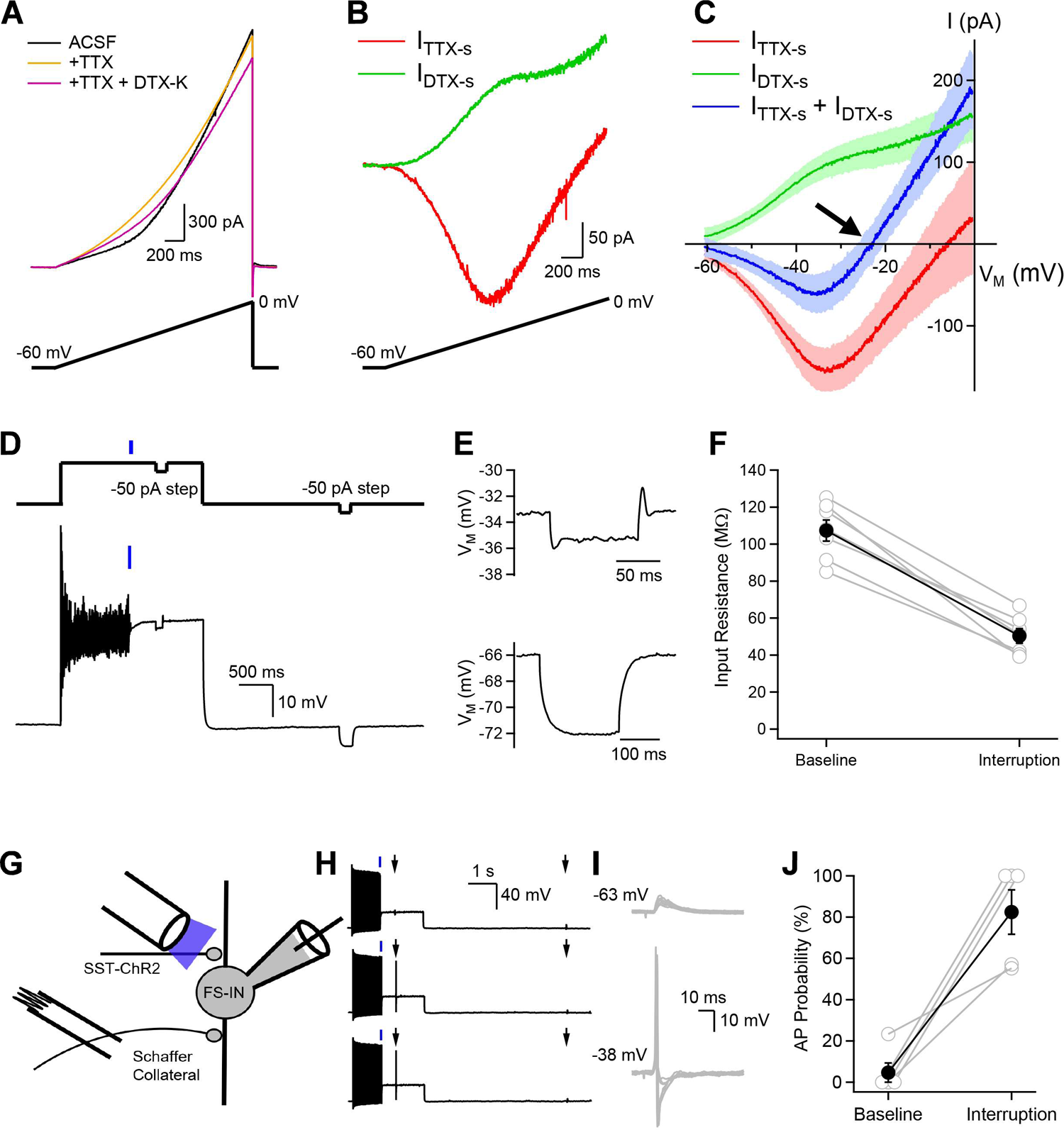
The interplay between K_v_1.1-current and Na^+^-current generates a stable point in membrane potential which results in a hyper responsive state. **A**, Voltage-clamp recordings from a PV-IN during ramp depolarization protocols. Data is shown in control (black), in presence of TTX (gold) and with both TTX and DTX-K present (purple). **B**, Arithmetic subtraction reveals the DTX-sensitive and the TTX-sensitive currents during the ramp depolarization protocol. **C**, Current plotted as a function of voltage for experiments presented in A and B. I_DTX-s_ and I_TTX-s_ were measured in the same neurons and shaded areas represent the standard error. **D**, Membrane potential dynamics during the firing interruption. Neurons were interrupted optogenetically, and brief hyperpolarizing current pulses of identical amplitude were applied during the interruption or at resting membrane potential. **E**, Membrane potential as a function of time for hyperpolarizing current injections delivered during the interruption (top) or at resting membrane potential (bottom) reveals drastically different dynamics. **F**, Input resistance measured at baseline and during the interruption from the same sweeps. **G**, Scheme showing the experimental design. Whole-cell current-clamp recordings were performed from PV-INs and neurons were optogenetically-interrupted. Schaffer collaterals stimulation was delivered during the interruption or at resting membrane potential by a stimulation electrode placed in CA3. **H**, Three consecutive sweeps showing that subthreshold EPSPs at rest become suprathreshold during the firing interruption. **I**, Changes in membrane potential evoked by Schaffer collaterals stimulation at resting membrane potential (top) or during the interruption (bottom). **J**, AP probability for stimuli delivered at resting membrane potential or during the interruption.

The involvement of inactivating K^+^ conductance harkens back to regulation of rhythmic AP firing (Connor and Stevens, 1971; Turrigiano et al., 1996; Goldberg et al., 2008; Khaliq and Bean, 2008). For example, I_A_ is de-inactivated by the afterhyperpolarization phase of an AP and slows down subsequent pacemaker depolarization in gastropod neurons (Connor and Stevens, 1971) and VTA dopaminergic neurons (Khaliq and Bean, 2008). Similarly, I_D_ temporally delays or negates firing in response to depolarizing current in cortical PV-INs (Goldberg et al., 2008; Campanac et al., 2013). In the present case, an inactivating K^+^ conductance (I_D_) also exerts a key braking action, but the initiating event is a synaptic input, and the outcome is a sustained cessation of ongoing firing rather than a graded delay in time-to-next-spike.

### The firing interruption maintains fast-spiking interneurons in a hyperresponsive state

Fast-spiking interneurons are known to be highly responsive to synaptic recruitment, more so than other elements of feedforward circuits (Fricker and Miles, 2000). This makes it interesting to determine how the interruption will alter PV-INs responsiveness to incoming synaptic inputs. The outcome is uncertain because opposing factors are at play in the PV-INs: on one hand, the elevation of intrinsic membrane conductance in the quiescent state and the lowered driving force for glutamate-induced current should dampen synaptic responsiveness. On the other hand, sodium channel activation should be enhanced at the depolarized membrane potential of the interruption, possibly promoting excitability.

EPSPs were evoked by electrical stimulation of Schaffer collateral inputs (Fig. 6G-I). The average EPSP evoked at resting membrane potential (−66.9 mV) had an amplitude of 6.53 ± 0.78 mV (n = 5), roughly 3-fold greater than a unitary EPSP of 2 mV (Miles, 1990; Fricker and Miles, 2000), as if the excitatory drive came from approximately 3 CA3 pyramidal cells. These EPSPs demonstrated a fast rise time (2.52 ± 0.4 ms, n = 5), consistent with previous findings (Fricker and Miles, 2000). The stimulation strength was adjusted to obtain mostly subthreshold EPSPs at resting membrane potential and spikes only rarely (AP probability = 4.67 ± 4.67 %, n = 5, Fig. 6H, I). We found that the same synaptic input from Schaffer collateral, evoked by a single electrical shock of fixed intensity (Fig. 6H, I), was much more likely to evoke an AP during the firing interruption (82.45 ± 10.75 %, n = 5; p < 0.01; Mann-Whitney U test; Fig. 6J). Thus, the firing interruption rendered the PV-IN super-responsive to incoming excitatory synaptic inputs.

### A minimal PV-IN model captures the interruption of firing and associated elevation in responsiveness

Interrupted neurons are in a shunted quiescent state but also hyperresponsive. Biophysical modeling of the experimental observations could add mechanistic insight into the interruption and possibly explain the hyperresponsiveness as well. As a first approximation, we used a model of the perisomatic region of the neuron to determine whether an interruption of firing could in principle arise from an interplay between intrinsic conductances and an IPSP-like hyperpolarization.

We assembled a minimal single-compartment model of a fast-spiking interneuron, incorporating transient Na^+^ current (I_Na_), delayed-rectifier K^+^ current (I_KDR_) and a small leak current (I_L_), based on Golomb et al. (2007) but supplemented by an inactivating K^+^ conductance (I_D_) previously described in CA1 hippocampal interneurons by Lien et al. (2002). With this combination of current components, the model reliably generated trains of APs in response to current injection (Fig. 7A). We then challenged the model with an incremental hyperpolarizing step with ramp recovery (triangular waveform, Fig. 7A1), identical to that in our experiments (Fig. 4B). Consistent with experimental observations, the model neuron’s firing was interrupted by such a protocol (Fig. 7A1). On the other hand, eliminating I_D_ from the model prevented the firing interruption (Fig. 7A2), an effect that persisted when depolarizing current amplitude was reduced to maintain the same evoked firing rate (Fig. 7A3). Moreover, the model replicated a telling aspect of the firing interruption: upon subsequent resumption of firing, the first AP was of smaller amplitude, consistent with the idea that sodium channels are partially inactivated (Fig. 7A1, inset). In addition, a rectangular hyperpolarizing pulse of varying duration (20 – 400 ms) failed to cause persistent interruption of the model neuron (Fig. 7A4), in line with experiment (Fig. 4F,G).

**Figure 7:**
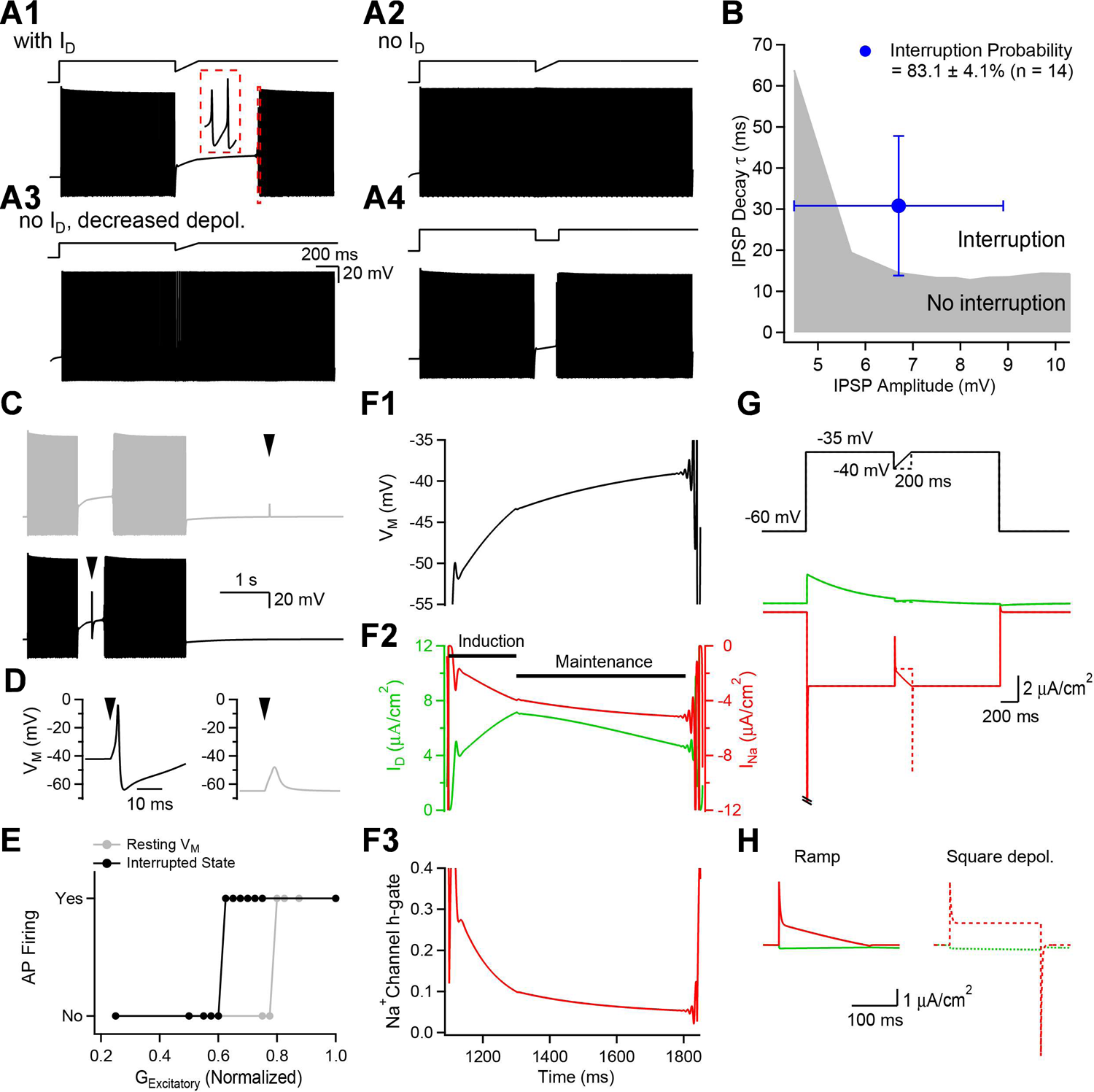
A single-compartment conductance-based model reproduces the core features of the firing interruption. **A**, Examples of firing returned by the model tested under different conditions. A hyperpolarizing pulse with a slow re-depolarization interrupts the model neuron in presence but not in absence of I_D_. Decreasing the depolarization to match the baseline firing frequency in presence of I_D_ could not rescue the interruption. In addition, a square hyperpolarizing pulse failed to interrupt firing (lower right). Inset in A shows the first two APs upon firing resumption. **B**, An inhibitory conductance in the model reliably interrupted firing. The model IPSP parameters required to interrupt firing were comparable to the properties of experimentally-measured IPSP sufficient to reliably generate the interruption (blue cross). In this comparison, the pre-interruption firing duration was kept constant (1 s) across experimental and modelling conditions. **C-E**, The model is hypersensitive to excitatory inputs during the firing interruption. An excitatory conductance (arrowhead) was inserted at resting membrane potential (gray) or during the interruption of firing (black). **D**, At resting membrane potential, the excitatory conductance is subthreshold while the same stimulus generates an AP during the interrupted state. **E**, Quantification of the excitatory strength required to generate an AP at resting membrane potential compared to that needed during the firing interruption. **F1**, Membrane potential as a function of time during the firing interruption. Note the slow and gradual depolarization observed during the interruption. **F2**, I_D_ and I_Na_ dynamics during the interruption. **F3**, Na^+^ channel inactivation variable (*h*-gate) as a function of time during the interruption segment reveals that PV-INs are increasingly accommodated. **G-H**, Interrogating the model in voltage-clamp with membrane potential dynamics known to interrupt neurons experimentally (hyperpolarizing step followed by a slow ramp re-depolarization, full line) or to cause resumption of their firing (square hyperpolarizing pulse, dotted line) simulates a firing resumption that is associated with a fast Na^+^ current.

Next, we aimed to explore the range of parameters allowing IPSPs to interrupt firing. An inhibitory conductance with an exponential decay was included in the model, and the conductance amplitude and decay time constant were systematically varied to determine pairs of parameters sufficient to interrupt firing (Fig. 7B). We observed that IPSPs over a broad range of amplitude could interrupt firing, but that smaller IPSPs required a longer decay time for the interruption to occur. Reassuringly, the IPSP parameters leading to a firing interruption in the model were generally similar to those measured in experiments (Fig. 7B). Also consistent with our experimental results (Fig. S2L-M), varying the E_Cl_ value in the model showed that hyperpolarizing and shunting inhibition (Vida et al., 2006) sufficed to interrupt firing, whereas depolarizing inhibition did not (Fig. S6A-B).

Our experimental findings indicate that interrupted neurons are in a quiescent but hyperresponsive state (Fig. 6). We next aimed to reconstruct this hyperresponsiveness. A compound EPSP-IPSP conductance sequence, simulating the experimentally observed outcome of Schaffer collaterals stimulation, was introduced at either resting membrane potential or during the interrupted state (Fig. 7C). Consistent with our experimental results, a subthreshold excitatory conductance at resting membrane potential became suprathreshold when imposed during the interrupted phase (Fig. 7D); this switch in excitability was seen over a broad range of amplitudes (Fig. 7E). Removing the IPSP conductance from the simulation, leaving only the EPSP, triggered the return of non-accommodating firing, indicating that the IPSP component had caused a renewal of the persistent interruption (Fig. S6C).

We took advantage of the model to look at the dynamic fluctuations in I_D_ and I_Na_ during the ramp decay and subsequent interruption (Fig. 7F1-F3 and Fig. S6D-E). Our results indicate that during the ramp decay (induction phase), I_D_ activates to hyperpolarize the neuron and therefore limits the speed of membrane potential re-depolarization, preventing AP firing (Fig. 7F2). During the maintenance phase, I_D_ inactivates gradually while I_Na_ gradually increases, resulting in a slow but steady membrane depolarization (Fig. 7F2). The gradual depolarization is paralleled by progressive Na^+^ channel inactivation (decreasing *h*), explaining why interrupted neurons are maintained in a non-firing condition (Fig. 7F3). Therefore, these results indicate that the interplay between I_D_ and I_Na_ not only initiates but also helps maintain the interruption of firing by first forcing accommodation and then by keeping the neuron in a depolarized yet quiescent state.

The above findings suggest that the pre-interruption state of I_D_ might influence the duration of the interruption. If I_D_ is more available when the interruption is initiated, the interruption may last longer. This could help reconcile the results from *in vivo* and *in vitro* experiments: interruptions are consistently longer *in vitro*, where longer and controlled depolarizing pulses are imposed. In both modelling and experiments, increasing the duration of pre-induction firing resulted in a progressive prolongation of the interruption (Fig. S7A-C). Examination of the underlying currents revealed that AP-evoked I_D_ gradually increased during firing episodes because of continuous I_D_ de-inactivation (Fig. S7B). I_D_ de-inactivation was driven by the large afterhyperpolarization (AHP) observed following every AP, and its build-up was attributable to slow I_D_ inactivation kinetics and the high firing frequency of PV-INs. In experiments, we observed that the AHP consistently hyperpolarized the membrane potential after allowing for an ohmic voltage drop across the series resistance during current injection (see Fig. S7 legends for details). The peak afterhyperpolarization in pooled data ranged from −84.2 ± 0.9 mV (1^st^ AP) to −76.8 ± 1.1 mV (20^th^ AP), consistently negative to resting levels (−65.6 ± 0.6 mV; n = 27; p 0.0001 for both comparisons). Thus, modeling and experiment converge to indicate that the duration of the firing interruption is influenced by I_D_ availability at the moment of incoming inhibition. This provides a mechanistic explanation for the lengthening of interruptions following increasingly prolonged firing. In both modelling and experiments, the interruption duration plateaued as pre-induction firing was prolonged (Fig. S7C). This can be attributed to a saturating degree of removal of I_D_ inactivation for longer firing episodes, evident in the model. The interruption duration varies with the extent of prior fast-spiking activity because such firing primes the neuron through I_D_ de-inactivation.

How is firing resumed following the interruption? We addressed this question by deconstructing two cases: rapid depolarization-induced firing and spontaneous firing resumption, both observed experimentally and in the model. First, an abrupt re-depolarization causes AP firing in experimental recording and modeling alike (Fig. 4F and Fig. 7A). We aimed to understand what interactions between I_D_ and I_Na_ support this firing recovery by voltage-clamping the model neuron (Fig. 7G) and applying changes in membrane potential known to trigger firing (offset of rectangular hyperpolarizing current pulse) or to interrupt the firing (IPSP-like ramp) (Fig. 7G).

We observed that a stepwise removal of hyperpolarization generated a large and fast I_Na_, a current which was not elicited by the ramp. The model predicts that a fast depolarization generates a Na^+^ current sufficiently fast and large to surmount the stable point in membrane potential, thus explaining the quiescent yet excitable state. This prediction was tested in experiments (Fig. S8A-D) by pharmacologically dissecting the currents during voltage changes identical to those observed during the firing interruption. Consistently, an abrupt depolarization generated a rapid and large I_Na_ which was not observed during an IPSP-like ramp. Altogether, these recordings confirm that the generation of a I_Na_ sufficiently large to escape the interruption of firing depends on the abruptness of the depolarizing stimulus. Second, we made further comparison between experiment and model for the case of spontaneous firing resumption. This was always preceded by membrane oscillations, gradually increasing in amplitude (17/17 neurons; Fig. S8E-G). Analyzing I_D_ and I_Na_ during a 35 ms oscillatory period right before firing resumption revealed that membrane potential-dependent oscillations emerged from a mismatch between the faster activation and inactivation kinetics of I_Na_ compared to I_D_, creating an instability in membrane potential, seen as a limit cycle in a phase plane plot (Fig. S8G).

### PV-IN interruption during elevated firing episodes powerfully disinhibits CA1 pyramidal cells

Our findings revealed that extended PV-INs firing episodes are highly prone to long-lasting interruptions. Extended PV-INs firing might occur physiologically under neuromodulatory influence because several neuromodulators, including oxytocin, can directly depolarize the membrane potential to drive rapid PV-INs firing (Owen et al., 2013; Tirko et al., 2018). This could render the cell susceptible to long-lasting interruptions, but on the other hand might also elevate firing probability leading to early termination of the silent period. At the level of downstream CA1 pyramidal cells, the consequences of such activity remain unknown but are particularly important given that pyramidal cell activity can be heavily influenced by the firing of even a single PV-IN (Cobb et al., 1995). We therefore aimed to understand the conditions favoring the occurrence of persistent firing interruptions and the direct consequences on CA1 pyramidal cell activity.

Application of the selective oxytocin receptor agonist (Thr^4^,Gly^7^)-oxytocin (TGOT) during generally subthreshold depolarization increased PV-IN firing rate tenfold (3.5 ± 1.79 Hz to 38.69 ± 12.24 Hz, n = 5, p < 0.05; Mann Whitney U test; Fig. 8A-B). In presence of TGOT, optogenetic stimulation interrupted firing with a high likelihood (97.66 ± 2.03%, n = 4), and silenced PV-INs persistently (821.8 ± 97.2 ms, n = 4; Fig. 8A-C). In presence of TGOT, PV-IN firing was resumed in subsets of trials (Fig. 8C), likely through the mechanisms described above. These results indicate that neuromodulatory enhancement of PV-IN activity can produce sustained firing episodes that are amenable to long-lasting firing interruptions. More generally, the interplay between local synaptic inhibition and neuromodulatory tone could provide a basis for abrupt switching of spiking in PV-INs.

**Figure 8:**
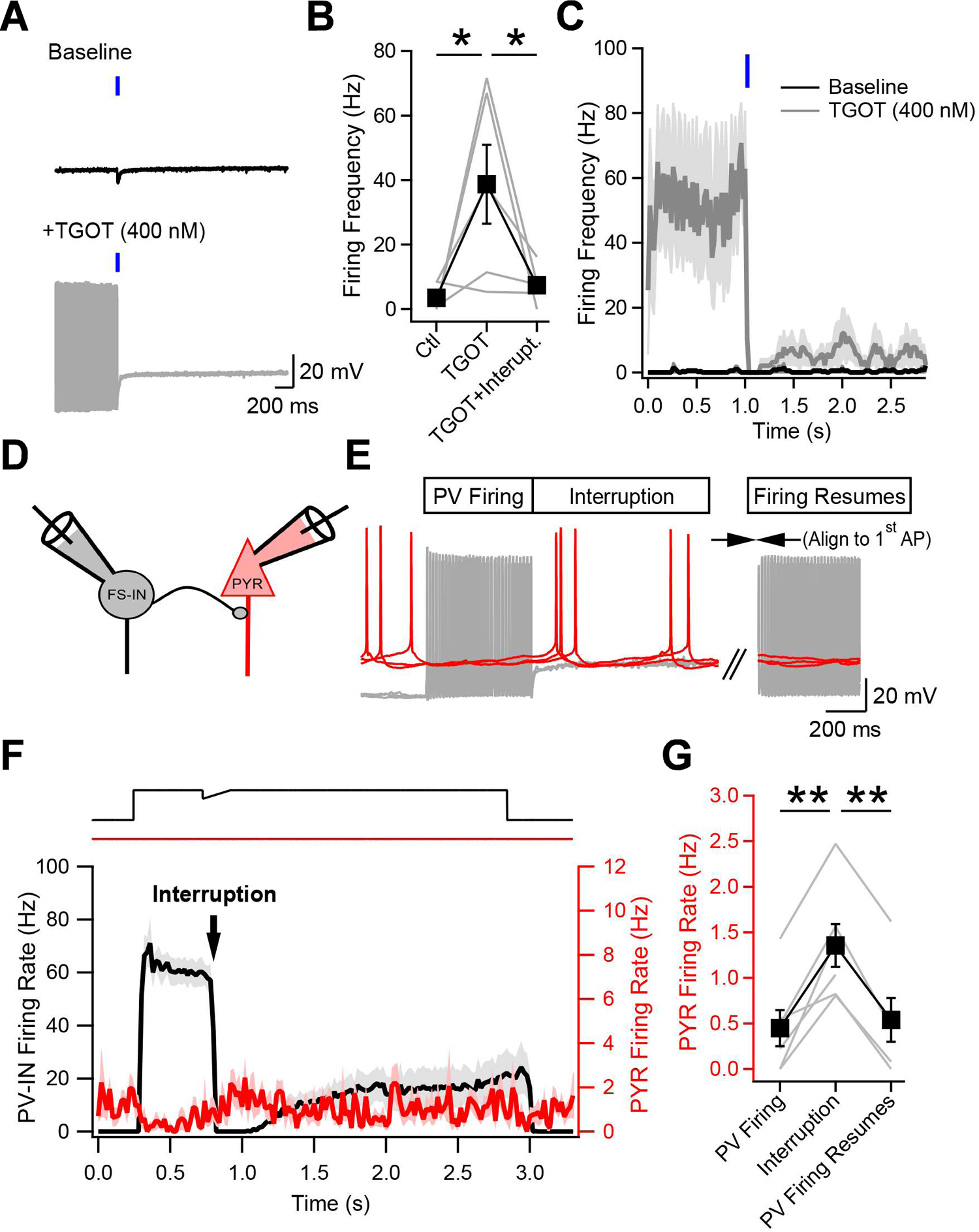
The firing interruption disinhibits CA1 pyramidal neurons. **A**, Current-clamp recording from a PV-IN at baseline (top) and following application of TGOT (bottom) to drive oxytocin receptor (OXTR) activation. At baseline, optogenetic activation of SST-INs (blue tick) causes only a brief GABAergic inhibitory response; after TGOT application drives AP firing, the same optogenetic activation of SST-INs causes persistent interruption of PV-IN firing. Thus, OXTR neuromodulation provides a platform for the interruption mechanism. 10 traces are shown overlayed for baseline and TGOT. **B**, Graph showing the pooled effect of TGOT and optogenetic activation of SST-INs on overall PV-IN firing (n = 5). **C**, Summary graph (n = 4) showing that optogenetically-evoked synaptic inhibition consistently and abruptly interrupts PV-INs driven to fire by OXTR activation. **D**, **E**, Paired whole-cell current-clamp recording from a PV-IN (black) synaptically connected to a CA1-PYR (red); 3 consecutive sweeps during the firing interruption induced by current injection as in F. **F**, Time course of average AP firing frequency in PV-INs (black) and CA1-PYRs for n = 6 neuron pairs. Interruption induced by IPSP-like hyperpolarization (top trace). Shaded areas correspond to the standard error. **G**, Summary graph showing the AP frequency recorded in the pyramidal cell for 500 ms windows measured during PV-IN firing, at the firing interruption onset (n = 6; ** p < 0.01) and following PV-IN firing resumption (n = 4; ** p< 0.01; 2 CA1-PYRs excluded because resumption of PV-IN firing was too rare to allow reliable assessment of pyramidal firing rate).

The observation that prolonged PV-IN silence periods could be observed during neuromodulator-driven firing episodes prompted us to investigate the consequence of PV-IN firing interruption and firing resumption on CA1 pyramidal cells. We aimed to determine how the interruption of firing in a single PV-IN directly impacts the activity of a downstream CA1-PYR target. Paired recordings were performed between PV-INs and deep CA1-PYR (Fig. 8D). We found that out of 65 attempts, 20 presynaptic PV-INs were synaptically connected to deep CA1 pyramidal cells as assessed by generation of IPSP in CA1-PYR by brief firing of PV-INs (30.8% connectivity rate). To assess the impact of the interruption, the CA1-PYR was slightly depolarized with current injection (30.5 ± 3.4 pA; n = 6) to allow tonic firing at 1 – 2 Hz (Fig. 8E), typical of CA1-PYR basal firing (Wiener et al., 1989; Czurko et al., 1999; Hirase et al., 1999). Meanwhile, the synaptically connected PV-IN was depolarized with steady current injection to drive firing and then suddenly interrupted with a mock IPSP. We observed that CA1-PYR firing was drastically decreased during bouts of PV-IN firing but returned to basal levels as soon as the PV-IN was silenced by a firing interruption (Fig. 8F). Indeed, pyramidal cell firing was elevated 3-fold during the interruption of firing (1.35 ± 0.24 Hz; n = 6) compared to that during PV-IN firing (0.45 ± 0.2 Hz; n = 6; p < 0.01; Fig. 8F,G). Remarkably, in trials where PV-INs firing subsequently recovered (interruption ceased), the CA1-PYR firing was decreased to similar levels as observed during initial PV-IN firing (0.54 ± 0.26 Hz; n = 4; p = 0.77; Fig. 8G) and was significantly lower than during the interrupted state (n = 4; p < 0.01). Thus, our results show directly that the firing interruption is a powerful disinhibitory mechanism for gating information flow. Interruption of firing in even a single presynaptic PV-IN suffices to elevate the firing activity of a downstream CA1-PYR.

## Discussion

Our experiments revealed that the apparently robust non-accommodating fast-spiking phenotype of hippocampal PV-INs is in fact a delicate state that can be toggled off by minimal synaptic inhibition, leading PV-INs to operate in a temporarily depolarized yet silent state. Once initiated, the persistent interruption of firing is a cell-autonomous condition that renders PV-INs quiescent yet hyperresponsive. In a circuit context, the persistent interruption of PV-INs firing not only removes their basal inhibition of CA1 pyramidal neurons, but also potentiates their responses to subsequent synaptic inputs, thus heightening feedforward inhibition-on-demand.

Our *in vivo* recordings displayed persistent silencing of PV-INs following optogenetically-induced synaptic inhibition that may share underpinnings with the persistent interruption of firing of PV-INs we studied *in vitro.* Insights into mechanism may help explain why the silences *in vivo* were generally briefer than persistent interruptions in acute slices. One factor is that interruption duration depends on recent firing history*—*briefer firing epochs preceding inhibition result in shorter interruptions afterwards*—*and the uncontrolled periods of PV-INs firing *in vivo* were greatly outlasted by the long high-frequency bursts we imposed for biophysical analysis *in vitro.* A second factor is that “interrupted” PV-INs would be hypersensitive; extrapolating to silenced PV-INs in freely moving animals, continual bombardment by synaptic inputs *in vivo* would often trigger early termination of the interrupted state. With these considerations in mind, our *in vivo* observations align with our highly controlled studies *in vitro* and *in silico*.

### Synaptic and intrinsic mechanisms controlling the interruption of firing

Neurons are endowed with intrinsic conductances that shape the impact of synaptic inputs in both duration and amplitude (Gulledge et al., 2005; Carter et al., 2012). The persistent interruption of firing is an extreme case of such amplification, wherein a brief IPSP de-inactivates I_D_, slows membrane re-depolarization, thereby partially inactivating I_Na_, and thus initiates the quiescent state.

Both pre- and postsynaptic dynamics contribute to the persistent interruption of PV-IN firing. Our paired recordings showed that GABA release evoked by a single AP from a PV- or SST-presynaptic partner can occasionally interrupt PV-IN firing, whereas brief bursts of inhibitory input trigger the interruption more reliably. At these synapses, the high release probability, large unitary currents and mild short-term depression during brief bursts of spikes (Bartos et al., 2001; Bartos et al., 2002; Hefft and Jonas, 2005; Bartos and Elgueta, 2012) are well-suited to interrupt PV-INs. This combination of features shapes a slow re-depolarizing ramp that is optimal to interrupt PV-INs as shown by direct current injection. Our observations indicate that any form of inhibition can interrupt PV-IN firing if it generates a hyperpolarization that is sufficiently large and slowly decaying.

After GABA_A_R conductance has decayed, the interruption of PV-IN firing is continued solely by intrinsic mechanisms. The non-accommodating fast-spiking pattern of PV-INs is supported by Na_V_1.1, Na_V_1.6 and K_V_3-family channels that enable rapid membrane depolarization and repolarization (Martina et al., 1998; Rudy and McBain, 2001; Lorincz and Nusser, 2008; Hu and Jonas, 2014). Although these currents are huge, the fast-spiking pattern they generate is prone to perturbation by the relatively modest currents provided by brief GABAergic input. The disparity sparks interest in the underlying biophysical mechanisms. Our reconstruction of the interruption splits it into two phases (Fig. 7F2). In the first (“induction”) phase, progressive I_D_ activation slows down the re-depolarization, partially inactivating I_Na_ and thus forestalling spiking. During the second (“maintenance”) phase, I_D_ inactivates to support a shallow but progressive depolarization, a delicate state of quiescence.

Both experiments and modelling converge in support of this scenario. Involvement of I_D_ was demonstrated with selective pharmacology and by corresponding omission of I_D_ in our computational model. In PV-INs, I_D_ was mediated by K_V_1.1- and K_V_1.2-containing channels which by themselves demonstrate little inactivation, therefore suggesting that beta subunits are incorporated and help shape the conductance dynamics. Given that K_V_1.1 is developmentally regulated in the hippocampus, the interruption of firing could be age-dependent (Pruss et al., 2010). The slow membrane re-depolarization progressively promotes I_Na_ inactivation, preventing a rebound spike. The maintenance phase is sustained by I_D_ inactivation and gradual I_Na_ activation, pitted against increasing outward current via I_KDR_ (Fig. S6D-E). This combination of current changes buffers the net current at a tiny inward value, driving a depolarization slow enough to keep the neuron quiescent.

The full impact of I_D_ on membrane trajectory depends on I_D_’s interplay with I_Na_, and I_KDR_. Together, these currents govern the interrupted state, elevating the membrane’s slope conductance compared to rest, yet rendering it hyperresponsive to depolarizing currents or excitatory synaptic inputs because of heightened Na^+^ channel activation. The elevated excitability is also manifested by emerging subthreshold membrane oscillations (Bracci et al., 2003; Golomb et al., 2007) whose growth gives way to the resumption of spontaneous firing, marking the end of the persistent interruption.

### Impact of persistent interruption of PV-INs firing on the CA1 hippocampal circuit

Intermittent silences would provide fast-spiking neurons more time to recover from the high metabolic demands they face (Cohen et al., 2018; Hu et al., 2018). Interruption of firing would also favor replenishment of presynaptic vesicle pools depleted by rapid firing (Kraushaar and Jonas, 2000; Park et al., 2021). Intermittency would give cell biological benefit to fast-spiking neurons whether they switched from fast firing to silent individually or collectively. Further advantages for network function might arise from concerted silencing of multiple PV-INs by an anatomically divergent presynaptic director. Ensemble silencing would engage a subset of PV-INs as a functional unit. Indeed, multiple place cells in CA1 can undergo coordination by concerted firing of their inhibitory afferents (Geiller et al., 2022). The monosynaptic inhibitory output from PV-INs provides further divergence, fanning out to contact >1500 pyramidal cells (Sik et al., 1995). Thus, mechanisms regulating the activity of PV-INs will be amplified anatomically, just as prolongation of GABA-triggered silencing of PV-INs from tens to hundreds of milliseconds would widen any impact of disinhibition.

Our paired recordings of PV-INs and CA1 pyramidal cells explored the consequences of the firing interruption on information processing in the CA1 circuit. Under conditions mimicking CA1-PYR resting state firing, synaptic inhibition by a single PV-IN decreased CA1-PYR firing rate by ∼3-fold. In turn, we demonstrated directly that shutting off this inhibition by an interruption of firing caused a rapid, powerful and consistent disinhibition of the local pyramidal neuron activity, an effect fully reversed by resumption of PV-INs firing. In parallel, we also showed that the interrupted state rendered PV-INs super-responsive to incoming inputs from the CA3 region, accentuating their potency as feedforward inhibitory elements (Buzsaki and Eidelberg, 1982; Fricker and Miles, 2000; Pouille and Scanziani, 2001), and possibly feedback inhibitory elements as well. Thus, feedforward inhibition is sensitized, dampening the net excitatory effect of input pathways. Altogether, the CA1 circuit will switch toward local information processing while veering away from receiving external inputs (Mizuseki et al., 2009; Mizuseki et al., 2012).

PV-INs strongly regulate CA1 population activity (Stark et al., 2013; Schlingloff et al., 2014), extending their influence in microcircuits. PV-INs, but not axo-axonic cells, are active during sharp wave-ripples (SPW-Rs), high-frequency oscillations associated with memory formation (Ylinen et al., 1995; Csicsvari et al., 1999; Klausberger et al., 2003; Klausberger and Somogyi, 2008; Viney et al., 2013). We speculate that regulating PV-INs firing by mechanisms like those found here could help control SPW-R duration, consistent with computational modeling of disinhibitory interactions during SPW-Rs (Evangelista et al., 2020). In turn, the duration of CA1 SPW-Rs strongly affects performance in hippocampally-based learning and memory tasks (Fernandez-Ruiz et al., 2019).

### Possible implications for disinhibition and pattern-switching in neocortical systems

In neocortex, *in vivo* studies have shown that PV-INs can experience intermittent bouts in a depolarized yet silent state close to AP threshold (Gentet et al., 2010; Yu et al., 2016). This raises the possibility that the persistent interruption of firing occurs outside the hippocampus and contributes more generally to *in vivo* regulation of PV-INs. In cortical areas, PV-INs are crucial in controlling neuronal network activity (Cardin et al., 2009; Sohal et al., 2009; Royer et al., 2012; Stark et al., 2013; Amilhon et al., 2015) and in regulating animal behavior (Donato et al., 2013; Kuhlman et al., 2013; McKenna et al., 2020). Disinhibition likely provides a permissive signal that allows input-selective integration by principal neurons (Lee et al., 2013; Karnani et al., 2016; Munoz et al., 2017; Turi et al., 2019). Inhibition of PV-INs is known to support learning and memory via downstream disinhibition of principal neurons (Letzkus et al., 2011; Wolff et al., 2014). Thus, more broadly beyond hippocampal CA1, the interruption of PV-IN firing and its net disinhibitory effect could participate in essential functions such as associative learning and spatially guided reward learning (Letzkus et al., 2011; Turi et al., 2019).

The persistent interruption of firing can be compared with forms of persistent network activity invoked to explain higher-order phenomena such as working memory and memory formation (Durstewitz et al., 2000; Egorov et al., 2002; Shu et al., 2003b). Networks have been found capable of maintaining an active condition in the absence of further external stimulation. The initiation of persistent activity can be cell-autonomous (Heyward et al., 2001; Egorov et al., 2002; Fuentealba et al., 2005; Loewenstein et al., 2005; Fransen et al., 2006; Tahvildari et al., 2007), sometimes reflecting integration of previous activity (Egorov et al., 2002; Loewenstein et al., 2005). In other cases, the maintenance of persistent activity requires continual neuromodulatory input (Egorov et al., 2002; Fransen et al., 2006; Tahvildari et al., 2007), engagement of other circuit elements (Shu et al., 2003b; Shu et al., 2003a), or participation of nearby astrocytes (Deemyad et al., 2018). In contrast, the persistent interruption of firing in PV-INs, while induced in a circuit context, is demonstrably sustained in a cell-autonomous manner. It is the first demonstration of switch-like changes in persistent firing activity initiated by a single presynaptic partner. Nonetheless, this simple flip-flopping between full-throated spiking or no firing could be an interactive building block of more complex circuit phenomena, incorporating neuromodulation, competing groups of neurons, non-neuronal partners and switching following integration of seconds-long trains of activity (Egorov et al., 2002; Fransen et al., 2006).

### Cooperation between persistent interruption of firing and slow neuromodulation

The interruption mechanism throws a new light on slowly acting neuromodulation. Oxytocin exemplifies agents that alter the intrinsic properties of PV-INs and drive them to fire rapidly and steadily. In this neuromodulatory setting, the firing interruption can relieve principal neurons from inhibition within milliseconds (Fig. 8D-G). The sharp transition would provide the kind of rapid disinhibitory switch invoked by Shen *et al*. to impose winner-take-all dynamics in a decision-making circuit (Shen et al., 2022). This disinhibitory scenario complements a distinct mechanism wherein spontaneous firing of PV-INs acts over many seconds to fatigue GABAergic synapses and thus weaken feedforward inhibition (Owen et al., 2013; Marlin et al., 2015). The common feature is an interplay between slow neuromodulators and fast GABAergic transmission that causes a net disinhibition of principal neurons. Such disinhibition could enable CA1 pyramidal cells to generate dendritic plateaus and potentially favor synaptic plasticity and place field formation (Magee and Grienberger, 2020).

## Material and Methods

### Animals

All experiments involving animals were approved by the Institutional Animal Care and Use Committee (IACUC) at New York University Langone Medical Center. For in vitro experiments, wild-type (C57BL/6) and transgenic mice (P17 – P30) of either sex were used indiscriminately in this study. For interneuron recordings in slices, homozygous Pv-Cre (Jackson Labs; Stock No. 008069) or Sst-IRES-Cre (Jackson Labs; Stock No. 013044) mice were crossed with homozygous Ai9 mice (Jackson Labs; Stock No. 007909) to generate *Pv;;Ai9* and *Sst;;Ai9* animals which demonstrated strong Td-Tomato expression in PV- or SST-expressing interneurons. For optogenetic stimulation of SST-expressing interneurons, homozygous Sst-IRES-Cre animals were crossed with homozygous Ai32 mice (Jackson Labs; Stock No. 024109). This cross resulted in offspring with channelrhodopsin-2(H134R) (abbreviated as ChR2 in figures) expression in SST-expressing interneurons (*Sst;;Ai32*).

### Acute hippocampal slice preparation

Acute hippocampal slices (300 μm) were prepared by deeply anesthetizing animals with isoflurane. The brain was rapidly extracted and placed in ice-cold slicing solution, containing (in mM): 185 sucrose, 25 NaHCO_3_, 2.5 KCl, 25 glucose, 1.25 NaH_2_PO_4_, 10 MgCl_2_, 0.5 CaCl_2_; pH 7.4, 330 mOsm. This solution was continuously oxygenated with a 95% O_2_ and 5% CO_2_ mixture. The brain was dissected, and slices were cut on a Leica VT1000 S Vibrating blade microtome. Slices were transferred to heated (32°C) slicing solution for 30 minutes, after which slices were transferred to oxygenated artificial cerebrospinal fluid (ACSF), containing (in mM): 125 NaCl, 25 NaHCO_3_, 2.5 KCl, 10 glucose, 2 CaCl_2_, 2 MgCl_2_; pH 7.4, 300 mOsm. Slices were left in this solution at room temperature for the duration of the experiment.

### *In vitro* electrophysiological recordings

Acute slices were transferred to a recording chamber and held under a nylon mesh. The preparation was continuously perfused with oxygenated ACSF (2 ml/min) at room temperature (20 ± 2°C, mean ± SD), unless otherwise indicated (Fig. S1H-I: 31.3 ± 0.9°C, mean ± SD). Recording electrodes were prepared from borosilicate filaments (TW150-4, World Precision Instruments) on a P-97 Sutter Instrument micropipette puller and had a resistance of 3 – 6 MΩ. For paired recordings, experiments were performed under an upright microscope (BX50WI, Olympus) equipped with a 40X objective. Whole-cell recordings were sequentially obtained by first bringing both recording electrodes (MP-285 micromanipulators, Sutter Instrument) close to targeted neurons and then forming giga-seals. For paired whole-cell electrophysiological recordings presented in Figs. 2 and 8, experiments were performed with a MultiClamp 700B amplifier and digitized at 10 kHz with a Digidata 1322A. Data was sent to a PC and acquired with the Clampex 9.2 software. All other electrophysiological recordings were performed with an upright microscope (BX61WI, Olympus) equipped with a 40X objective. The electrophysiological signal was amplified with an Axopatch 200B, digitized at 10 kHz (Digidata 1322A) and recorded on a PC equipped with the Clampex 8.2 software. The intracellular solution contained (in mM): 130 K-gluconate, 10 HEPES, 2 MgCl_2_.6H_2_O, 2 Mg_2_ATP, 0.3 NaGTP, 7 Na_2_-Phosphocreatine, 0.6 EGTA, 5 KCl; pH 7.2 and 295 mOsm. Under these conditions, the total intracellular [Cl^-^] was 9 mM and the theoretical Cl^-^ reversal potential was −69 mV. In experiments with elevated intracellular [Cl^-^] reported in Fig. S2L-M, the intracellular solution contained (in mM): 121.5 K-gluconate, 10 HEPES, 2 MgCl_2_.6H_2_O, 2 Mg_2_ATP, 0.3 NaGTP, 7 Na_2_-Phosphocreatine, 0.6 EGTA, 13.5 KCl; pH 7.2 and 295 mOsm. Under these conditions, the total intracellular [Cl^-^] was 17.5 mM and the theoretical Cl^-^ reversal potential was −52 mV. Only cells with a series resistance below 25.7 MΩ were included. Series resistance was 18.12 ± 0.72 MΩ for current-clamp recordings presented in Fig. 1. Series resistance in the voltage-clamp recordings presented in Fig. 6A-C was 19.71 ± 1.68 MΩ (n = 8) and was not compensated. Schaffer collaterals were stimulated by positioning a tungsten electrode connected to a stimulus isolator (A360, World Precision Instruments) in the stratum radiatum of the CA3 region. Photostimulation of SST-INs was performed with 470 nm light from a light-emitting diode (LED) delivered to the slice with an optical fiber. A TTL signal was sent from the digitizer to an LED controller for precisely timed stimulation (WT&T inc.). For voltage-clamp recordings, neurons were held at the indicated potential in the figures. The liquid junction potential was not corrected. The following pharmacological reagents were used in this study: tetrodotoxin (1 μM, Sigma), bicuculline (10 μM, Sigma), CGP-55845 (2 μM, Tocris) dendrotoxin-K (50 nM, Alomone), dendrotoxin-I (50 nM, Alomone), K-Conotoxin RIIIK (200 nM, Alomone), Agitoxin-2 (10 nM, Alomone), TGOT ((ThrJ,GlyJ)-oxytocin, 400 nM, Bachem).

### *In vivo* electrophysiological recordings and optogenetic stimulation

All experiments were approved by the Institutional Animal Care and Use Committee (IACUC) at New York University Medical Center. *Sst;;Ai32* mice (n = 2; 28-35 gr, 4-6 months old; from Ssttm2.1(cre)Zjh/J, Jax stock number: 013044 and B6.Cg-Gt(ROSA)26Sortm32(CAG-COP4*H134R/EYFP)Hze/J, Jax stock number: 024109) were implanted with 64-site silicon probes (NeuroNexus A5×12-16-Buz-lin-5mm-100-200-160-177) in dorsal CA1 (AP 2.0 mm, ML 1.6 mm, DL 1.1 mm). Ground and reference wires were implanted in the skull above the cerebellum, and a grounded copper mesh hat was constructed shielding the probes. Probes were mounted on microdrives that were advanced to pyramidal layer over the course of 5-8 days after surgery. A 100 µm fiber optic was attached to the silicon probe (Valero et al., 2021). The back end of the fiber was coupled to a laser diode (450 nm blue, Osram Inc.). Animals were allowed to recover for at least one-week prior to recording. Mice were housed under standard conditions in the animal facility and kept on a 12 h reverse light/dark cycle. Electrophysiological data were acquired using an Intan RHD2000 system (Intan Technologies LLC) digitized with 30 kHz rate. For optogenetic tagging of Sst-expressing neurons, blue laser light (450 nm, Osram Inc) pulses were delivered. The maximum light power at the tip of the optic fiber was 1 to 4 mW. 20, 50 and 100 ms light pulses were delivered (n = 500 - 1000 times at each duration at 400 ± 200 ms random intervals).

### Biocytin revelation, neuronal tracing, and anatomical classification

Neurons were passively filled with biocytin in the whole-cell configuration. Following recordings, the pipette was carefully retracted, and the acute slice was placed in a petri dish between filter papers. Slices were fixed overnight with 4% PFA in PBS. Biocytin was revealed by treating the slices with Triton (1%) and incubating overnight in an Alexa-633 conjugated streptavidin (1:200, ThermoFisher Scientific). The following day, slices were mounted on microscope slides with ProLong Gold (ThermoFisher Scientific). Images were acquired on a Zeiss confocal system (Axo Imager.Z2). Anatomical tracings were performed in Neurolucida 360 (2.70.1, MBF Bioscience) on a personal computer.

For anatomical classification, the axonal length in the dendritic layers (strata oriens and radiatum) and in the somatic layer (stratum pyramidale) were quantified in Neurolucida. For each cell, axonal length was measured using Neurolucida 360. The axonal length in the somatic or dendritic layers were then normalized to the total axonal length for each cell. Using this dataset, K-means clustering analysis in Python was used to cluster interneurons in two groups.

### Stereotaxic injections

For stereotaxic surgeries, mice were anesthetized with isofluorane (2%–5%) and secured in a stereotaxic apparatus (Kopf). Glass pipettes (Drummond Scientific) were formed using a P-2000 puller (Sutter Instrument) and were characterized by a long taper and 10-20 *μ*m diameter tips. Pipettes were back-filled with mineral oil (Fisher Scientific) before being loaded with pertussis toxin (Sigma P7208) and positioned over the lateral ventricle (coordinates relative to bregma, in mm: 0.25 lateral, 0.3 anterior, −3 ventral). A small drill hole was made in the skull to allow for pipette insertion. 1 – 2 *μ*L of 0.1 g/L pertussis toxin were injected unilaterally into the ventricle. Experiments were performed 24 – 72 hours following injection. Throughout the surgery, body temperature, breathing and heart rate were monitored. Saline was administered subcutaneously (s.c) to maintain hydration and the animal was monitored post-operationally for signs of distress and discomfort. Buprenorphine (0.1 mg/kg, s.c) was given for analgesia. No major adverse effects of the surgery or pertussis toxin injection were observed.

### Immunohistochemistry

For localization of K_V_1.1 in PV-IN, 20 μm thick hippocampal slices from Pv-Ai9 animals were prepared on a cryostat (CM3050 S, Leica). Slices were treated with a K_v_1.1 recombinant rabbit monoclonal antibody (SN66-06, ThermoFisher Scientific) overnight and with an Alexa-488 conjugated secondary antibody for two hours on the following day. Images were acquired on a Zeiss confocal system (Axo Imager.Z2). Pv-Ai9-expressing interneurons were considered positive for K_v_1.1 if the Alexa-488 fluorescence intensity at the soma was two standard deviations above the surrounding background.

### Computational modeling

A conductance-based fast-firing interneuron model was conceived from previously published data obtained in ModelDB (senselab.med.yale.edu/modeldb/) (Golomb et al., 2007). The model was implemented in NEURON (version 7.7). The model consisted of a single cylindrical compartment with a diameter of 10 μm and a length of 10 μm. Axial resistance was set to 100 Ωcm, membrane capacitance was set to 1 μF/cm^2^ and the leak conductance was set to g_pas_ = 0.0001 S/cm^2^ with a reversal potential of −65 mV. The model contained a Na^+^ conductance (Na_t_; reversal potential: 50 mV; g_Na_ = 0.1125 S/cm^2^) and a delayed-rectifying K^+^ conductance (K_dr_; reversal potential: −90 mV; g_Kdr_ = 0.225 S/cm^2^) (Golomb et al., 2007) as well as an inactivating K^+^ conductance (K_D_) (Lien et al., 2002). These conductances were modeled using the Hodgkin-Huxley formalism. Parameters of Na_t_ and K_dr_ were left unchanged. The maximum conductance G_D_ of the inactivating K^+^ conductance was empirically determined based on the firing frequency measured experimentally before (77 Hz) and after DTX treatment (90 Hz) and set to 0.01 S/cm^2^. Temperature during simulations was set to 24°C. Excitatory and inhibitory synaptic conductances were modeled with a double-exponential time course of onset and decay. Excitatory currents had rise and decay times of 0.2 ms and 2 ms, a maximum conductance of 0.3 nS, and a reversal potential of 0 mV. Inhibitory currents had rise and decay times of 1 ms and 50 ms, a maximum conductance of 0.6 nS, and a reversal potential of −65 mV. Decay time and maximum conductance of inhibitory synapses were systematically varied to generate Fig. 7B. Simulations were performed with a step size of 0.025 ms. Simulations were performed on a personal computer in the NEURON interface controlled by Python and simulated traces were analyzed in Igor Pro 6.37 (Wavemetrics).

### Electrophysiological data analysis

*In vitro* electrophysiological data was analyzed in Clampfit 10.3 (Molecular Devices) and in Igor Pro 6.37 (Wavemetrics). The likelihood of observing a firing interruption was obtained by dividing the number of sweeps showing a successful interruption by the total number of acquired sweeps. An interruption was deemed successful if the silence period exceeded the IPSP duration. The IPSP duration was measured from its initiation to 95% recovery. The interruption duration was measured as the time from the IPSP onset to time of the first AP after firing resumption. For graphs representing the AP frequency as a function of time, the timing of the AP was determined at its peak amplitude and the data was binned in 20 ms width.

For *in vivo* electrophysiological data analysis, spike sorting was performed semi-automatically with KiloSort 47 (https://github.com/cortex-lab/KiloSort), using our own pipeline KilosortWrapper (a wrapper for KiloSort, DOI; https://github.com/brendonw1/KilosortWrapper). This was followed by manual adjustment of the waveform clusters using the software Phy2 (https://github.com/kwikteam/phy) and plugins for Phy designed in the laboratory (https://github.com/petersenpeter/phy-plugins). The following parameters were used for the Kilosort clustering: ops.Nfilt: 6 * numberChannels; ops.nt0: 64; ops.whitening: ‘full’; ops.nSkipCov: 1; ops.whiteningRange: 64; ops.criterionNoiseChannels: 0.00001; ops.Nrank: 3; ops.nfullpasses: 6; ops.maxFR: 20000; ops.fshigh: 300; ops.ntbuff: 64; ops.scaleproc: 200; ops.Th: [4 10 10]; ops.lam: [5 20 20]; ops.nannealpasses: 4; ops.momentum: 1./[20 800]; ops.shuffle_clusters: 1.

Unit clustering generated three separable groups (Fig. 3B) based on their autocorrelograms, waveform characteristics and firing rate. Putative pyramidal cells, narrow-waveform interneurons and wide-waveform interneurons were tentatively separated based by these three clusters (Valero et al., 2022). Definitive cell identity was assigned after inspection of all features, assisted by monosynaptic excitatory and inhibitory interactions between simultaneously recorded, well-isolated units and optogenetic responses. Units were defined as optically tagged using a p value cutoff of 10^-3^ (Valero et al., 2021).

### Statistical Treatment

For *in vitro* electrophysiological data, Shapiro-Wilk test was performed to test for normality of data distribution. For normally distributed data, a paired or unpaired Student’s t-test was performed to evaluate statistical significance. For non-normally distributed data, a Mann-Whitney U test was used where indicated. Pearson rank correlation was used to evaluate correlation between parameters in Figs. S1F and 4D-E. A two-way ANOVA was used to evaluate statistical significance in Fig. S7C. Experimental groups were deemed significantly different if p < 0.05. Statistical tests were performed in Clampfit 10.3 (Molecular Devices) and in Python. Statistical significance is reported on figures as follows: * p < 0.05, ** p < 0.01, *** p < 0.001.

Statistical analyses for *in vivo* electrophysiological data were performed blinded or did not require manual scoring and were performed with standard MATLAB functions. No specific analysis was used to estimate minimal population sample and the number of animals, trials, and recorded cells were similar to those employed in previous works (Valero et al., 2021; Valero et al., 2022). Unless otherwise noted, for all tests, non-parametric two-tailed Wilcoxon’s paired signed-rank test and Kruskal-Wallis one-way analysis of variance were used. When parametric tests were used, the data satisfied the criteria for normality (Kolmogorov–Smirnov test) and equality of variance (Bartlett’s test for equal variance). For multiple comparisons, Tukey’s honesty post hoc test was employed and the corrected *p < 0.05, **p < 0.01, ***p < 0.001 are indicated, two-sided. Boxplots represent median and 25th/75th percentiles and their whiskers the data range. In some of the plots, outlier values are not shown for clarity of presentation, but all data points and animal were always included in the statistical analysis. The exact number of replications for each experiment is detailed in the text and figures.

## Acknowledgements

We thank Dr. Michael A. Long for valuable comments throughout the execution of this project, Dr. Guoling Tian for technical support and maintenance of animal colonies, and all Tsien lab members for comments and discussions. SC was supported by a senior biomedical postdoctoral fellowship from the Charles H. Revson Foundation, a postdoctoral fellowship from the Fonds de Recherche en Santé Québec and a K99/R00 Pathway to Independence Award from NIMH (1K99MH126157-01). MV was supported by postdoctoral fellowships from the European Molecular Biology Organization (EMBO ALTF 1161-2017) and Human Frontiers Science Program (LT0000717/2018). RE was supported by a Research Fellowship from the Deutsche Forschungsgemeinschaft (EG 401/1-1). SBL was supported by postdoctoral fellowships from the NIA (1T32AG052909-01A1) and from the Alzheimer’s Association (AARF-21-852397). Work in GB’s lab was supported by the following grants: NIH MH107396, NS 090583, NSF PIRE (grant no. 1545858), U19 NS107616. RWT received grants from the NINDS (1U19NS107616-02), NIDA (R01 DA040484-04) and NIMH (R01 MH071739-15).

## Author contribution

SC and RWT conceived the project. SC performed *in vitro* electrophysiological experiments and analyzed the data. MV and GB designed, performed, and analyzed *in vivo* recordings. ERN and MH analyzed neuronal anatomy. MH and SBL performed immunohistochemistry experiments. KE performed pertussis toxin injections. RE and SC designed the single-compartment model, and SC ran simulations. SC and RWT wrote the manuscript with inputs from all authors.

## Code availability

All custom code for analysis of *in vivo* data is freely available on the Buzsáki Laboratory repository (https://github.com/buzsakilab/buzcode) and MV repository (https://github.com/valegarman/HippoCookBook).

## Data availability

The data from this study is available on request from SC or RWT. The *in vivo* electrophysiological data used in this study is publicly available on the Buzsaki Lab Databank, https://buzsakilab.com/wp/public-data/.

