## Supplemental Text and Figures for "Brief synaptic inhibition persistently interrupts firing of fast-spiking interneurons"

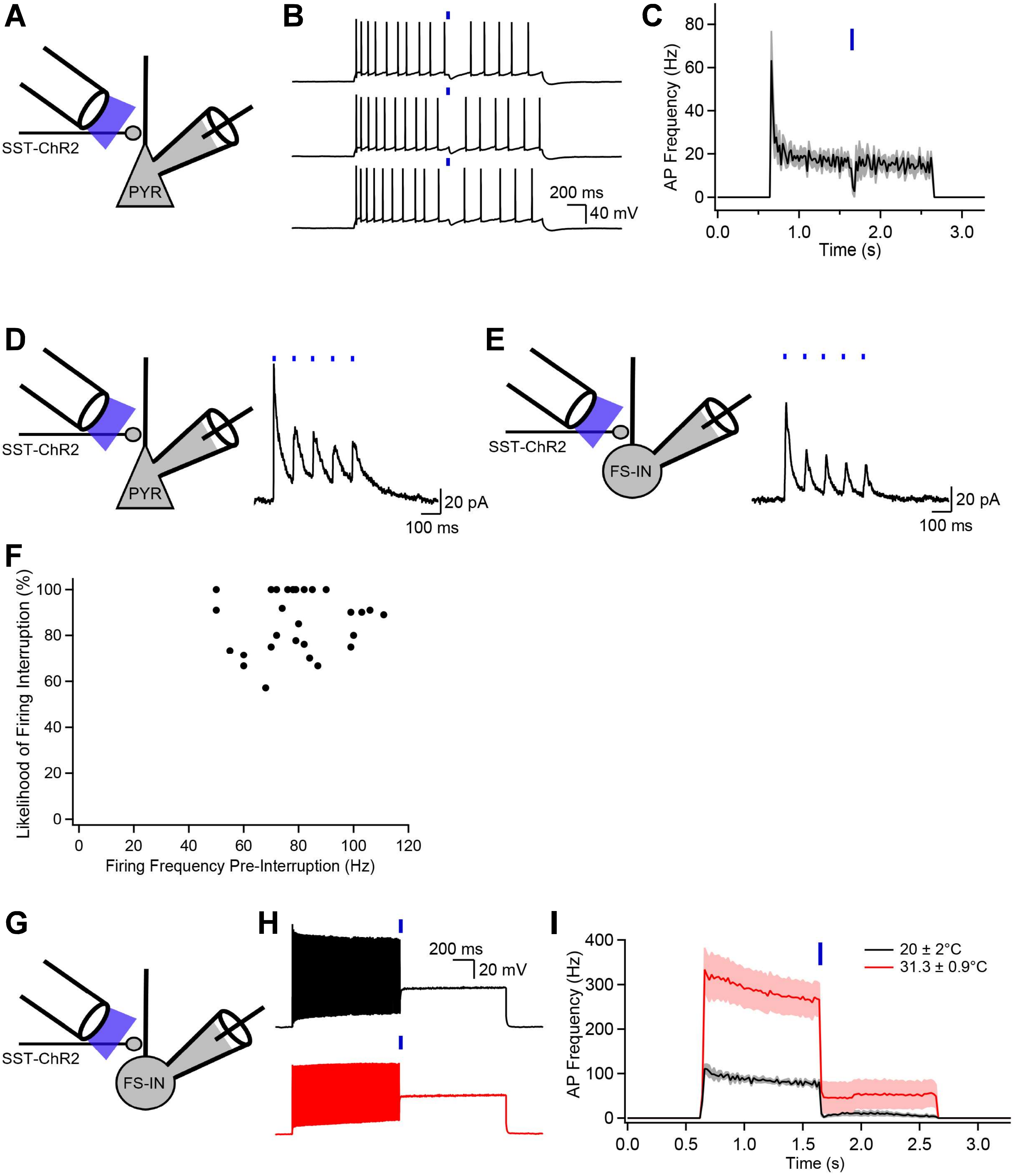


**Supplementary Figure 1**

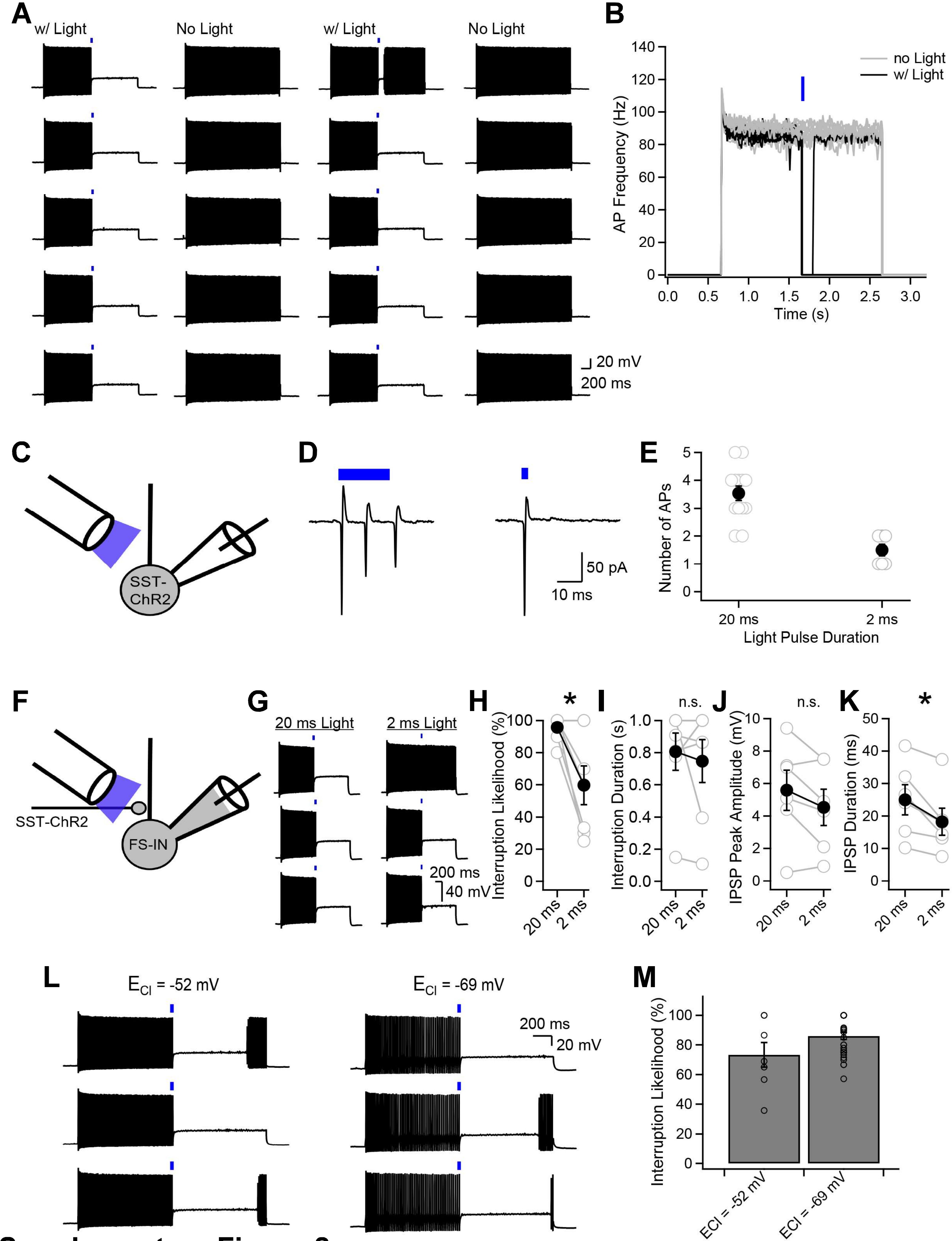


**Supplementary Figure 2**

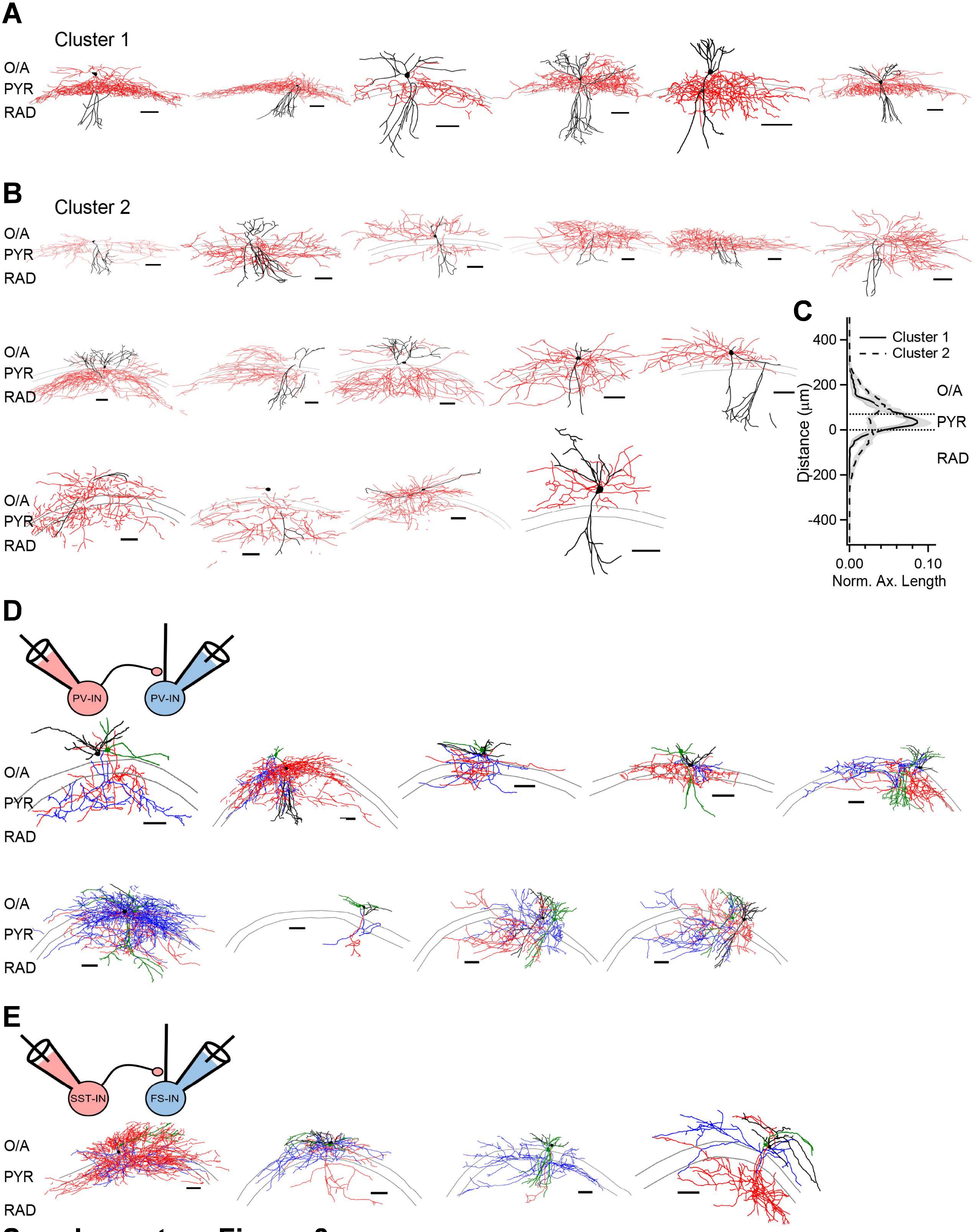


### Supplementary Figure 3

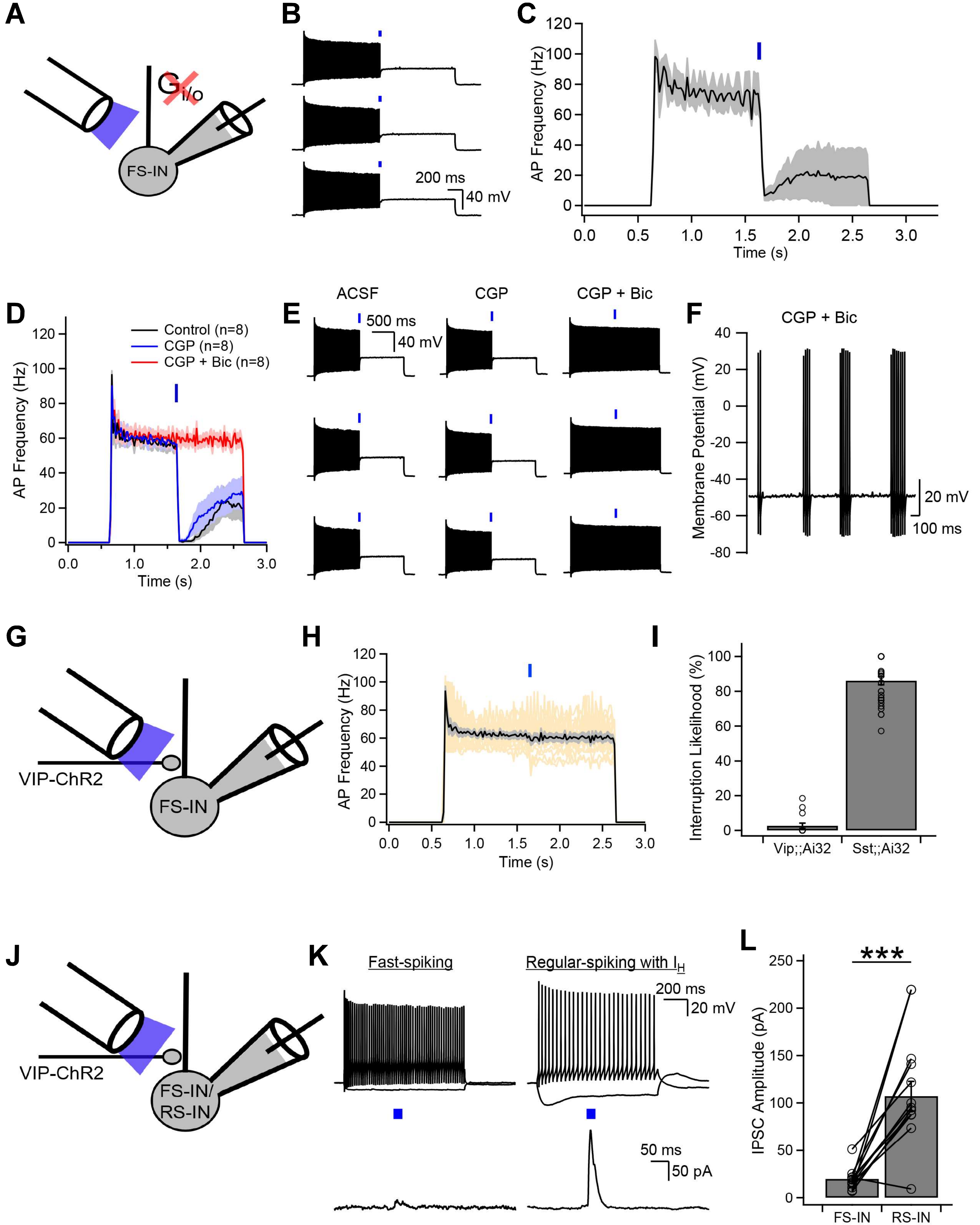


**Supplementary Figure 4**

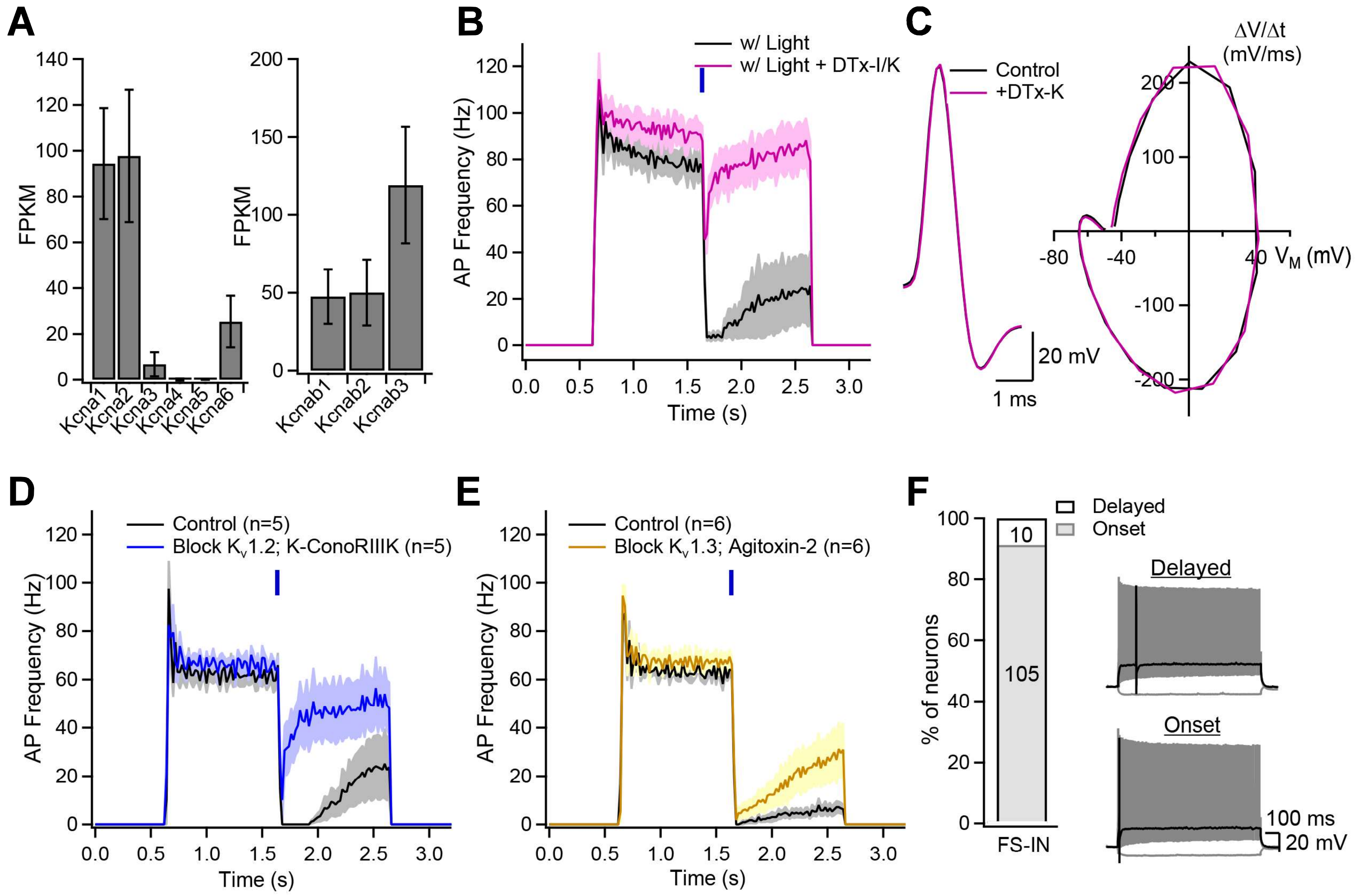


### Supplementary Figure 5

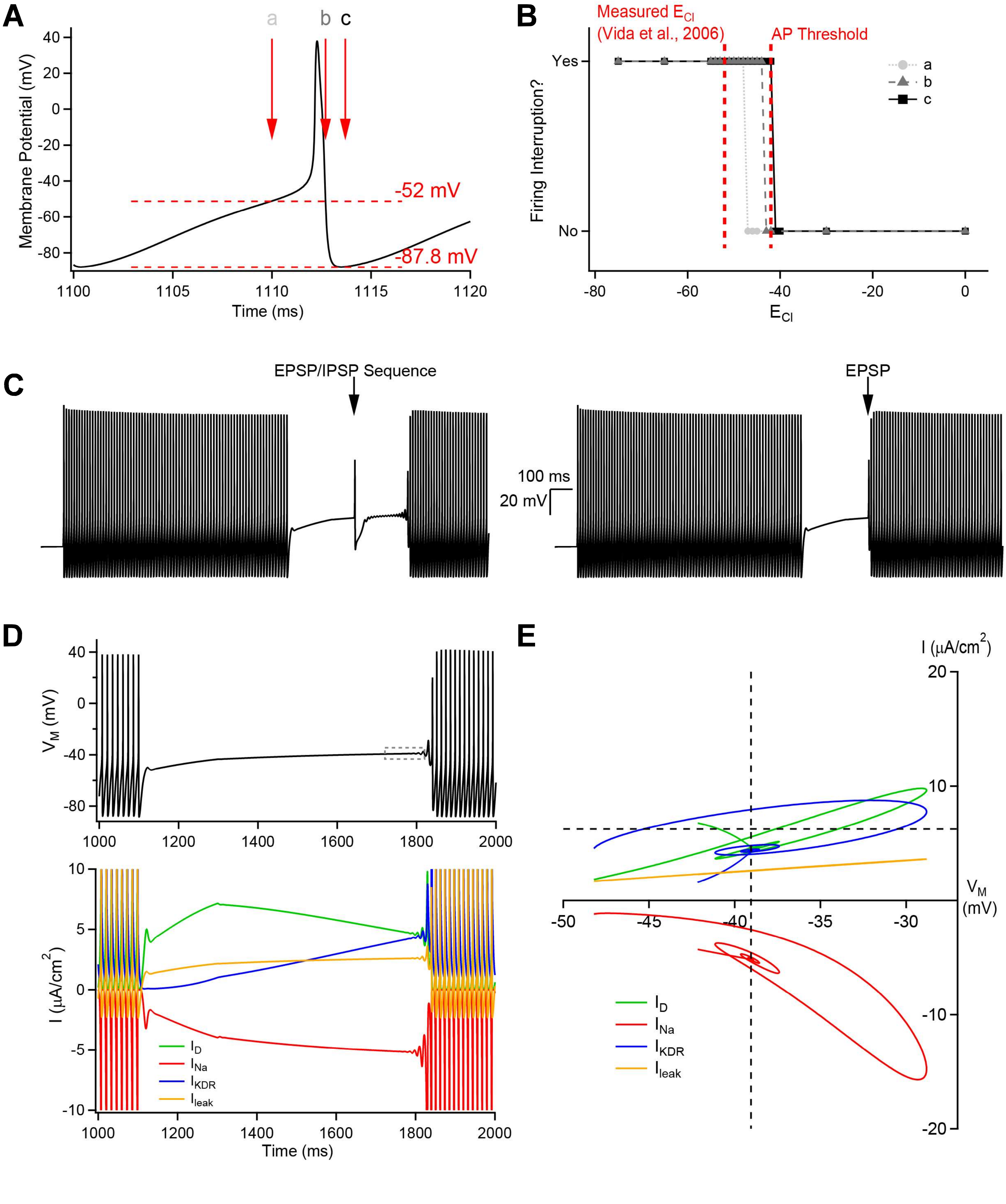


### Supplementary Figure 6

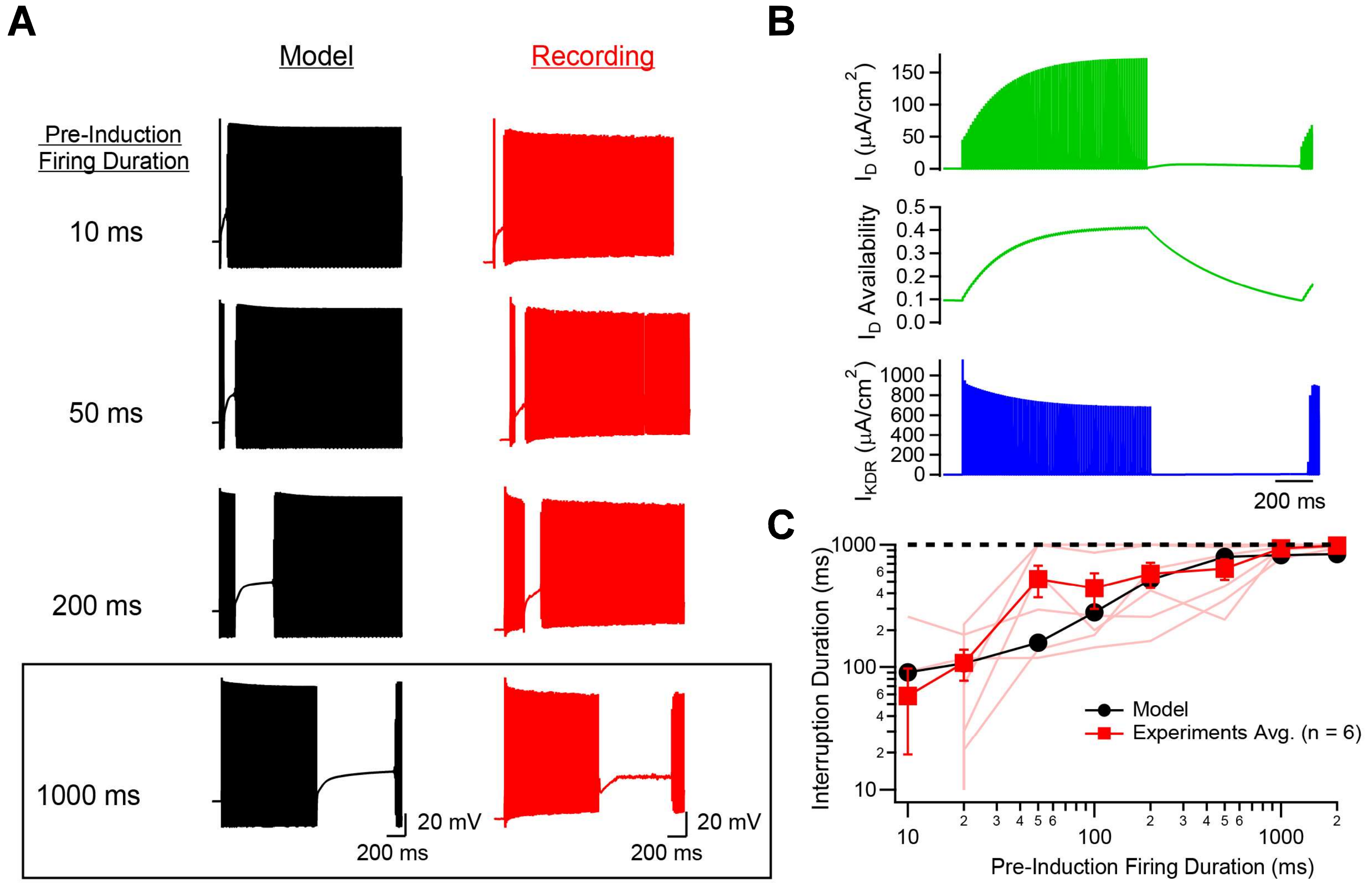


**Supplementary Figure 7**

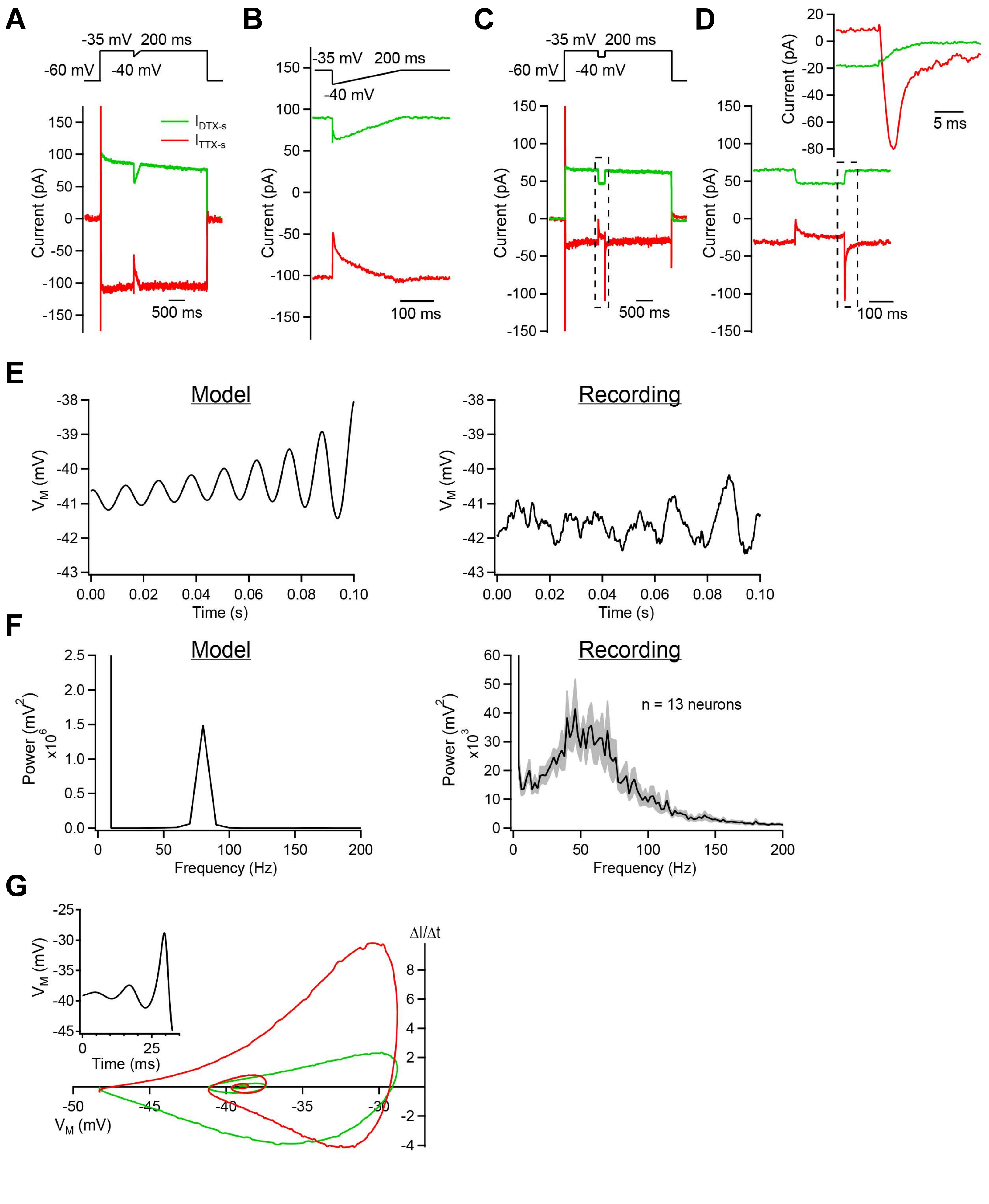


### Supplementary Figure 8

#### Supplemental Information

##### Supplementary Figure Legends

**Supplementary Figure 1: Optogenetic stimulation does not interrupt CA1 pyramidal cell firing and characterization of the firing interruption**

**A**, Experimental scheme. **B**, Three consecutive sweeps showing that optogenetic stimulation of SST-expressing GABAergic afferents does not interrupt CA1-PYR, which rapidly resume their spiking activity. **C**, AP frequency as a function of time for the 7 CA1-PYR recorded and tested with this protocol. **D**, **E**, Voltage-clamp recordings from CA1-PYR (**D**) or from PV-INs (**E**) show similar IPSCs during repetitive optogenetic stimulation (5 pulses delivered at 20 Hz). **F**, Likelihood of firing interruption as a function of the firing frequency prior to the interruption. **G**, Recording scheme. **H**, Current-clamp recordings showing the firing interruption for a neuron at room temperature and for an experiment performed at physiological temperature. **I**, Summary graph of neurons recorded at 20 ± 2°C (mean ± SD, black) and 31.3 ± 0.9°C (mean ± SD, red). In addition, we tested a range of temperature (33°C – 37°C) at which the interruption of firing was observed. 3/3 neurons tested demonstrated the interruption of firing at 33.5 ± 0.4°C (mean ± SD). 1/1 neuron tested demonstrated the interruption of firing at both 35°C and 37°C. Note the high baseline firing rate at physiological temperature. Consequently, the firing frequency is higher at physiological temperature upon firing resumption.

**Supplementary Figure 2: Presynaptic and postsynaptic determinants for the firing interruption**

**A**, Consecutive epochs of firing interruption interleaved with depolarization of same current amplitude but in absence of optogenetic stimulation. The firing interruption is highly reliable in response to optogenetic stimulation. **B**, Summary graph showing the firing frequency as a function of time for the data presented in (A), including the 10 trials with optogenetic stimulation and the 10 control trials without illumination. **C**, Cell-attached recordings were performed from SST-expressing interneurons. **D**, Exemplar cell-attached recordings during 20 ms or 2 ms optogenetic stimulation. **E**, Graph showing the number of APs as a function of optogenetic stimulation duration. **F**, Recording scheme. **G**, Three consecutive sweeps showing firing interruptions induced by 20 ms or 2 ms optogenetic stimulation. **H**, The likelihood of observing an interruption of firing is significantly higher for longer optogenetic stimulation. **I**, Upon initiation, the duration of the firing interruption is independent of the optogenetic stimulation duration, likely pointing to a postsynaptic mechanism. **J**, IPSP peak amplitude and duration (**K**) for 20 ms and 2 ms light pulses. **L**, Representative examples of recordings performed with two intracellular $Cl^-$ concentrations and summary data for all recordings (**M**).

**Supplementary Figure 3: Anatomical reconstructions of interneurons**

Neurolucida reconstructions of recovered fast-spiking interneurons reported in this studies. Dendrites are in black, and axons in red. All scale bars represent 100 µm. Cluster analysis based on axonal distribution revealed two population of neurons with an axon innervating the perisomatic (**A**) or the dendritic regions (**B**). **C**, Normalized axon length as a function of distance from the pyramidal cell – stratum radiatum intersection. The axon of cluster 1 neurons is found within close distance of stratum pyramidale while the axon of cluster 2 interneurons is mostly found in the dendritic layers. Shaded areas represent the standard error. **D**, **E**, Neurolucida reconstructions of recovered interneurons from paired-recording experiments. The presynaptic neuron is shown in black and its axon is shown in red. The postsynaptic neuron is shown in green and its axon is in blue. The last pair shown in panel D was reciprocally connected and is shown twice in both color combinations.

**Supplementary Figure 4 – The firing interruption is insensitive to $GABA_B$ receptor antagonism, does not require $G_{i/o}$ signaling and is not induced by VIP-INs activation**

**A**, Recordings were performed from animals injected with PTx for 24 – 48 hours. **B**, **C**, Firing interruptions were reliably induced in neurons recorded from PTx-treated animals, indicating that the interruption does not rely on G-protein dependent mechanism(s). **D**, **E**, Example traces and summary graph showing that the $GABA_B$R antagonist CGP has no effect on the firing interruption but that subsequent bicuculline application fully prevented the interruption in those neurons. **F**, PV-INs can still spontaneously stutter in presence of CGP and bicuculline. **G**, Recording scheme. **H**, AP frequency as a function of time showing that VIP-INs activation does not interrupt PV-INs firing. Individual cells are shown in yellow and their average is shown in black with shaded area representing the standard error (n = 15 neurons). **I**, Summary graph showing that optogenetic activation of VIP-INs is largely insufficient to interrupt PV-INs (n = 15 neurons) compared to optogenetic activation of SST-INs (n = 29; same data presented in Fig.

1D). **J**, Recording scheme. **K**, **L**, In sequential recordings from neighboring neurons, photoactivation of VIP-INs resulted in large amplitude IPSCs in regular-spiking INs with anodal breaks but significantly smaller IPSCs in neighboring fast-spiking INs.

**Supplementary Figure 5 – Dissection of $K_v$1-family subunits supporting the firing interruption**

**A**, Expression of $Kcna_{1\text{-}6}$ and $Kcnab_{1\text{-}3}$ subunits in PV-INs. **B**, Dendrotoxin treatment affects baseline firing rate and interruption likelihood. Summary graph showing the AP frequency as a function of time in control and after DTX treatment. The current injection was kept constant, which resulted in a higher baseline firing rate. Thus, the current injection was reduced to match control data and avoid potential confounding effects of increased AP generation likelihood (data presented in Fig. 5G). **C**, AP before (black) and after (purple) DTX-I/K treatment. The AP take-off potential was significantly decreased (control: -44.52 ± 0.82 mV; DTX-I/K: -45.52 ± 0.67 mV; $p < 0.05$; n = 8) and no significant difference were observed in the maximal dV/dt (control: 225.8 ± 5.53 mV/ms; DTX-I/K: 232.19 ± 7.47 mV/ms; p = 0.12; n = 8) and in the AP peak amplitude (control: 108.4 ± 1.8 mV; DTX-I/K: 108.4 ± 2.3 mV; p = 0.98; n = 8). **D**, **E**, Summary graphs showing the AP frequency as a function of time for experiments testing the effect of $K_v$1.2 and $K_v$1.3 blockade on the firing interruption. **F**, Number of INs presenting onset- and a delayed-firing phenotypes.

**Supplementary Figure 6: Current dynamics during the interruption of firing in the single-compartment model**

**A**, Effect of setting the $Cl^-$ reversal potential ($E_{Cl}$) to different values in the model. **B**, Outcomes of multiple simulations with varying $E_{Cl}$, showing the binary outcomes of firing interruptions being successful (Yes) or nonsuccessful (No). For all $E_{Cl}$ values below the AP take-off potential leading to shunting or hyperpolarizing inhibition, the injection of an inhibitory postsynaptic potential results in a firing interruption. **C**, While the EPSP/IPSP sequence results in the firing of a single AP, an EPSP by itself caused the return of non-accommodating firing. **D**, AP firing in the model is interrupted by a step-ramp waveform and activity of the four currents in the model during the segment of activity shown above. **E**, Current-voltage relationships of the four currents denoted, following the interruption and just before spontaneous firing resumption. The segment of time depicted is denoted with a dashed box in (D).

**Supplementary Figure 7: The firing interruption duration depends on past firing history**

**A**, Simulations showing that longer pre-induction bursts are associated with increased firing interruption duration. Experiments confirming the dependence of the interruption duration on the pre-induction firing. Presented traces are from raw data that do not account for the ohmic voltage drop across the series resistance during current injection. Once this voltage drop is factored in, the AHP consistently attains a peak hyperpolarization negative to the resting membrane potential (see Results section), in conformity with simulations indicating a key role for the AHP in driving de-inactivation of $I_D$ and induction of the interruption of firing. The horizontal box indicates the 1000 ms pre-induction firing duration used as an exemplar in (B). Interruptions were induced by an inhibitory conductance in the model and a ramp re-depolarization in experiments (as in Fig. 4B). **B**, Dissection of the membrane current dynamics underlying the 1000 ms pre-induction firing duration. **C**, Summary graph showing a similar dependence of the interruption duration on the pre-induction firing duration in simulations and experiments (n = 6 neurons; p = 0.18; two-way ANOVA).

**Supplementary Figure 8: A damped oscillation and fast $Na^+$ currents are associated with firing resumption**

Voltage-clamp recordings were performed to dissect the current interactions underlying firing resumption. **A**, $I_D$ and $I_{Na}$ dynamics during a membrane potential waveform experimentally observed during interruption. **B**, Under this condition, $I_D$ matches the kinetics of $I_{Na}$ during the ramp re-depolarization. **C**, $I_D$ and $I_{Na}$ during a square hyperpolarizing pulse protocol that resumes firing. **D**, The fast re-depolarization generates a fast and large $Na^+$ current as predicted by the model. **E**, A growing subthreshold oscillation is observed in the model and in experiments before spontaneous firing resumption. **F**, Power spectrum analysis of the subthreshold oscillation in the model and in pooled experiments. The shaded area for the recordings show the standard error. **G**, Incremental changes in $I_D$ and $I_{Na}$ plotted as a function of $V_M$ for the recording segment shown in the inset highlights faster activation and inactivation kinetics of $I_{Na}$ relative to $I_D$ as a mechanistic explanation for the subthreshold membrane oscillations. Inset shows subthreshold membrane oscillations during the interruption used for the investigation of current dynamics.
